# A fitness trade-off explains the early fate of yeast aneuploids with chromosome gains

**DOI:** 10.1101/2022.07.07.499182

**Authors:** Simone Pompei, Marco Cosentino Lagomarsino

**Affiliations:** IFOM ETS - The AIRC Institute of Molecular Oncology, via Adamello 16, Milan, Italy; Dipartimento di Fisica, Universitá degli Studi di Milano, via Celoria 16, Milan, Italy; INFN sezione di Milano, via Celoria 16, Milan, Italy

## Abstract

The early development of aneuploidy from an accidental chromosome missegregation shows contrasting effects. On the one hand, it is associated to significant cellular stress and decreased fitness. On the other hand, it often carries a beneficial effect and provides a quick (but typically transient) solution to external stress. These apparently controversial trends emerge in several experimental contexts, particularly in the presence of duplicated chromosomes. However, we lack a mathematical evolutionary modeling framework that comprehensively captures these trends from the mutational dynamics and the trade-offs involved in the early stages of aneuploidy. Here, focusing on chromosome gains, we address this point by introducing a fitness model where a fitness cost of chromosome duplications is contrasted by a fitness advantage from the dosage of specific genes. The model successfully captures the experimentally measured probability of emergence of extra chromosomes in a laboratory evolution setup. Additionally, using phenotypic data collected in rich media, we explored the fitness landscape, finding evidence supporting the existence of a per-gene cost of extra chromosomes. Finally, we show that the substitution dynamics of our model, evaluated in the empirical fitness landscape, explains the relative abundance of duplicated chromosomes observed in yeast population genomics data. These findings lay a firm framework for the understanding of the establishment of newly duplicated chromosomes, providing testable quantitative predictions for future observations.

## Introduction

Aneuploidy, a deviation from the normal chromosome number, is a form of large-scale genomic variation, involving changes both at the genotypic and at the phenotypic level, and one of the hallmarks of cancer [1]. In cancer genomes, aneuploidy correlates with important genomic changes, such as TP53 mutation and expression of proliferation genes [2] and drug resistance mutations leading to treatment failure [3, 4]. Drugs that disrupt mitotic progression, called antimitotic drugs [5, 6], are widely used for cancer treatment. These drugs cause chromosome missegregation, large genetic rearrangements and aneuploidy. In *S. cerevisiae* models (hereafter referred to as yeast), perturbed gene expression due to extra chromosomes can cause stress resulting from the proteome-wide stoichiometric imbalance of protein levels [7, 8]. Moreover, aneuploidy was shown to cause global changes of mRNA and protein expression and to possibly confer condition-dependents fitness advantage [9–11]. Finally, in yeast, the ploidy levels were shown to influence the emergence of mutator strains [12], to alter the affect of genetic mutations [13], to influence the phenotypic attributes of hybrid individuals [14], to impact the speed of adaptation [15, 16], to act on transcriptional silencing [17] and on pleiotropic effects [18].

The evolutionary dynamics leading to the emergence of aneuploidy is typically investigated with yeast models, because they can be manipulated with advanced genetics and cell- and molecular-biology methods, and hence be used to create isogenic backgrounds that only differ from each other by chromosome ploidy number, offering a direct point of comparison between euploid and aneuploid strains. Moreover, yeast models can be easily investigated with laboratory-evolution experiments, thanks to their short replication time [6, 9, 19, 20].

Several experiments in the last decades have raised apparently controversial evidence for the evolutionary role of aneuploidy [10, 21]. While aneuploidy carries significant cellular stress and decreased fitness [22], measured for example by a reduction of growth rate, it has also been shown to carry a beneficial effect that provides a quick, transient solution to external stress [20, 21]. This quick solution often emerges faster, hence, more frequently, than other evolutionary routes. Intriguingly, cultured human cells show the same contrasting trends: aneuploid cancer cells lines show a reduction of growth rate [23–25], but specific patterns of aneuploidy, particularly in the presence of extra chromosomes, confer a beneficial effect in specific adverse conditions [26, 27].

In the case of yeast, the literature offers extensive phenotypic data, for example growth curves of aneuploids in several environmental conditions as well as in laboratory-evolution experiments [6, 9, 10, 19–21], offering the opportunity to test for unifying trends. The available modelling studies presented so far have focused on the effect of aneuploidy on cell growth and physiology [7, 28–31]. Two interesting recent studies [28, 30] proposed a stochastic model of evolution similar to the classic Fisher’s geometrical model [32] to describe the fitness landscape of a set of engineered aneuploid strains in different stress environments. This model explains the observed correlation between the degree of phenotypic variation and the degree of overall growth suppression, measured in [9]. However, this model, and all the approaches presented so far [7, 28–31], are limited by a static description of the genetic and phenotipic architecture of aneuploids, failing to provide a description of the mutational dynamics leading to its emergence.

Here, we develop a theoretical framework to describe such mutational dynamics and to address the cost-benefit trade-offs in early aneuploids. We introduce a fitness model where a fitness cost proportional to the number of genes in the duplicated chromosome is counterbalanced by a fitness advantage resulting from the dosage increase of specific genes. Our approach builds on the so called “mutation bias” framework [33–35], a class of evolutionary models used to investigate the role of fast mutational processes in directing evolution, in a scenario where evolutionary routes emerging with a high mutation rate are in evolutionary competition with alternative mutational targets generated with a lower rate, but able to confer higher fitness advantage. Our model makes quantitative predictions that capture the dynamics leading to the emergence of aneuploidy. As we will describe in detail, the model captures the probability of the emergence of extra chromosomes in experimental setups [20],and correctly predicts the observed outcomes for the emergence of aneuploidy. We then make use of phenotypic data to isolate the main features contributing to the fitness landscape of aneuploids with extra chromosomes, and show that the dynamics of our model in this landscape captures the relative abundance of aneuploidies observed in population genomics data.

## Results

### Model and parameters

We develop and analyze an evolutionary model to describe the emergence of aneuploidy carrying extra chromosomes. Fig. 1A describes the key model ingredients. The model considers a population consisting of euploid individuals, which is exposed to an external stress causing a decrease of their growth rate. Individuals in the population can respond to the stress by increasing the expression of a specific target gene, gaining a beneficial effect quantified by the selection coefficient (*σ*_*b*_ *>* 0). Individuals can gain fitness by two alternative evolutionary routes: (i) by increasing the target gene expression (for example with mutations on the promoter binding site) or improving its functionality via a set of point mutations (on coding regions adapting protein function), occurring at a total rate *μ*_*m*_ or (ii) via missegregation events, taking place at a higher rate (*μ*_*a*_ *> μ*_*m*_) and resulting in the emergence of aneuploid individuals carrying extra chromosomes. Note that route (i) could require several point mutations (modeled here as a one-step process), but the per-base mutation rate is a lower bound for *μ*_*m*_. Aneuploids with extra chromosomes are less fit than euploids, because the duplication of the non-target genes in the extra chromosome determines a global fitness cost (*σ*_*c*_ *>* 0). Hence, the selection coefficient of aneuploids (*σ*_*b*_ − *σ*_*c*_) is lower than that of euploid mutants (*σ*_*b*_) and euploid mutants, although generated at a lower rate, have a higher fixation probability than aneuploids (*ϕ*_*m*_ *> ϕ*_*a*_). Double mutants (individuals carrying both aneuploidy and point mutations) are produced at a rate corresponding to the product of the rates (*μ*_*m*_ *× μ*_*a*_) and therefore are very rare and can be neglected. We also assume for simplicity that all mutations other than missegregations and the target point mutations do not contribute to the adaptive dynamics in response to the external stress, hence are neglected. Under these assumptions, the model reduces to the competition between two possible beneficial mutations, aneuploidy *vs* a local mutation.

**Figure 1:**
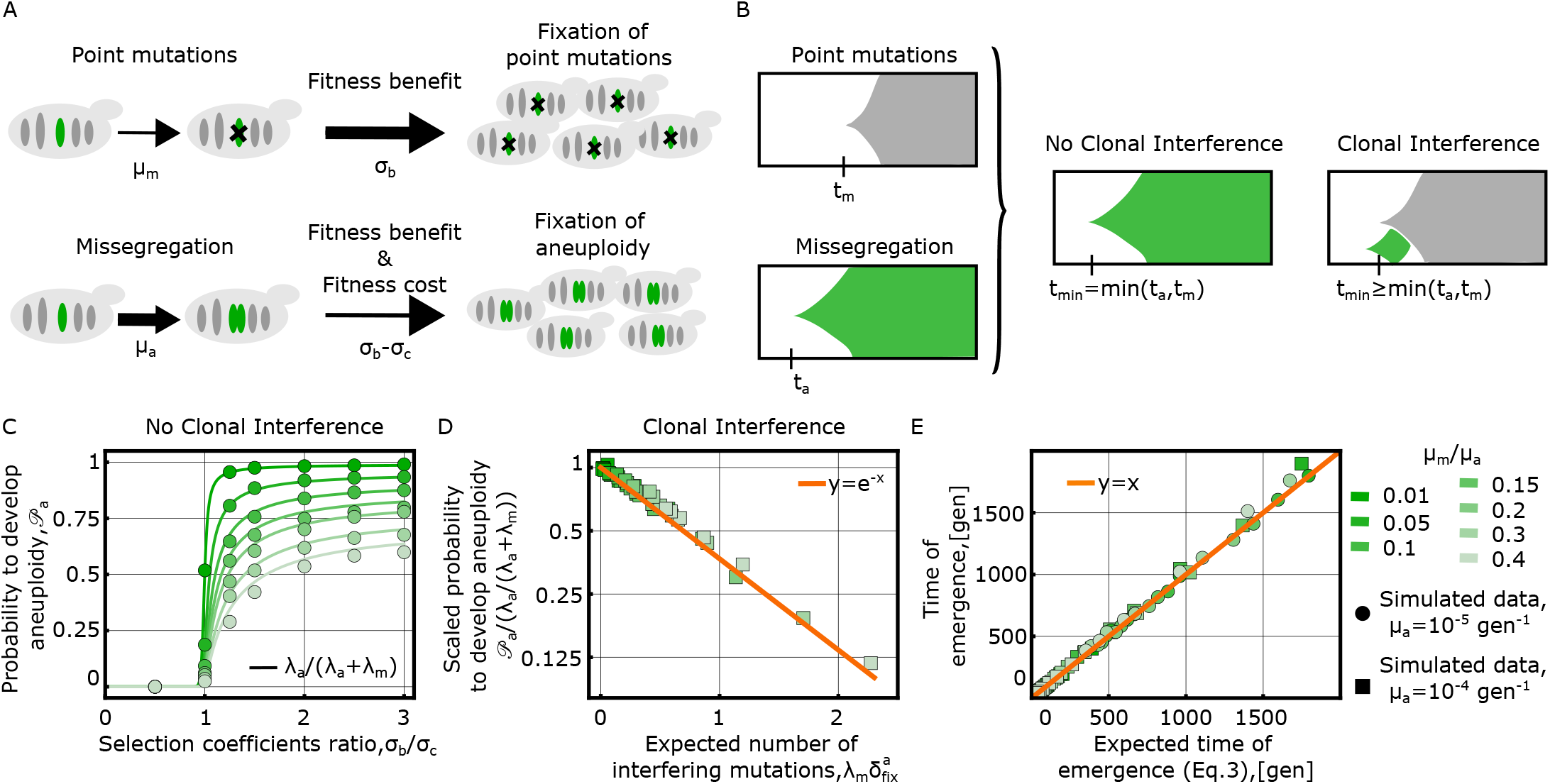
A trade-off between fitness cost and fitness benefit explains the population dynamics of early aneuploids with extra chromosomes. A: Schematic illustration of the evolutionary model, which considers two alternative mutational routes to cope with an external stress requiring the increase of the expression of a specific target gene. Individuals in an evolving population can increase the target gene expression via point mutations, taking place at rate (*μ*_*m*_), or via duplication of the target chromosome, at a rate related to missegregation. This second evolutionary route takes place at a higher rate (*μ*_*a*_ *> μ*_*m*_) and generates aneuploids. Aneuploids with extra chromosomes pay a fitness cost, because they carry the duplication of non-target genes in the extra copy of the chromosome containing the target gene. Hence, euploids, although emerging at a lower rate, have a higher fixation probability than aneuploids *ϕ*_*m*_ *> ϕ*_*a*_. B: The dynamics of the two evolutionary routes, here schematically represented with Müller plots,which are used to illustrate the succession of genotypes in an evolutionary process (the horizontal axis shows time, while vertical axis represents relative abundances of genotypes. Each genotype is shown with shaded areas of different colors, and originates in an arbitrary clone placed in the middle of its parent area), is characterized by the time of emergence of the successful mutant (the mutant whose descendants will eventually take over the population). In the model, both evolutionary routes are attainable and only the fastest of the two mutants, (i.e. the one that will generate a mutation able to overcome the genetic drift and reach fixation), emerging at a time *t*_min_ = min(*t*_*a*_, *t*_*m*_), will reach fixation. Clonal interference, occurring for example when a euploid mutant emerges during the fixation dynamics of an aneuploid individual, will prevent the fixation of aneuploidy and effectively delay the emergence of the successful mutant (*t*_min_ ≥ min(*t*_*a*_, *t*_*m*_)). C: Fixation probability of aneuploids with extra chromosomes in a regime with no clonal interference 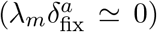, plotted as a function of the non-dimensional ratios *σ*_*b*_*/σ*_*c*_ (x axis) and *μ*_*m*_*/μ*_*a*_ (color coded). Results of simulations (circles) are compared to analytical calculations (solid lines). D-E: Collapse plots of simulated data of the model in the clonal interference regime 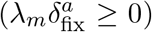 validate the analytical results for the fixation probability of aneuploids with extra chromosomes (Eq. 1, shown in panel D) and for the emergence time of the successful mutant (Eq. 3, shown in panel E). See Material and Methods for details on the numerical simulations of the model. Additional model parameters used for the data shown in panels B,C,D: *N* = 1000, *σ*_*c*_*N* = 50.

Of note, the benefit of the aneuploidy is modelled for simplicity as having the same selection coefficient (*σ*_*b*_) as the euploid beneficial mutation. However there could be many possible mutations that alter gene expression to different degrees (with a distribution of effect sizes) of those effects may differ compared to a gene duplication that presumably doubles expression. This scenario translates into a lower selection coefficient for the euploid mutant. The reduced selection coefficient would result in a lower fixation probability *ϕ*_*m*_ and in a lower fixation rate. This same effect is effectively described by a reduction of the mutation rate 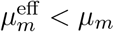, while keeping the same fixation probability (and hence, the same selection coefficient). In other words, for the fixation dynamics, it is equivalent to have a lower selection coefficient or to have a lower mutation rate, allowing to map on our model scenarios where point mutations of euploid individuals alter the expression of the target gene to a lower degree than a full chromosome duplication (this point is further explained in SI Appendix).

### Evolutionary dynamics

Our question concerns the conditions in which chromosomal duplications emerge first. Accordingly, we investigate the “early stage” population defined by the point in time when one of the two mutants, the euploid with point mutations or the aneuploid mutant carrying extra chromosomes, becomes fixed in the population (i.e., reaches an intra-population frequency ≃ 1) for the first time. This dynamics is described by fixation rates, which are given by the product of the mutation rates, the effective population size *N*, and the fixation rates: *λ*_*m*_ = *μ*_*m*_*Nϕ*_*m*_(*σ*_*m*_, *N*) for the euploid mutant and *λ*_*a*_ = *μ*_*a*_*Nϕ*_*a*_(*σ*_*a*_, *N*) for the aneuploid one. The fixation probabilities depend on the selection coefficients (*σ*_*a*_ ≡ *σ*_*b*_ − *σ*_*c*_ for the aneuploid, and *σ*_*m*_ ≡ *σ*_*b*_ for the euploid mutant) and on the effective population size *N*, as given by Kimura’s formula *ϕ*(*σ, N*) = (1 − *e*^−2*σ*^)*/*(1 − *e*^−2*σN*^) [36].

### Analytical expression of the probability to develop aneuploidy with extra chromosomes

In order to characterize the onset of the fastest variant, we focus on the waiting times for the emergence of a successful mutant, defined as the mutant that will eventually reach fixation. The two times, denoted as *t*_*a*_ and *t*_*m*_ for the aneuploid carrying extra chromosomes and the euploid mutant respectively, are stochastic variables. Since these mutations emerge at a constant rate, the probability distribution of the waiting times is exponential *t*_*a,m*_ ∼ *Exp* (*λ*_*a,m*_), hence, their expected values are equal to the inverse of the fixation rates (*τ*_*a,m*_ ≡ *⟨t*_*a,m*_*⟩* = 1*/λ*_*a,m*_).

The statistics of the fastest emerging mutant can be described by the difference of the two times, *t*_diff_ ≡ *t*_*a*_ − *t*_*m*_, whose probability density has an analytical expression (see SI appendix). In particular, the problem of computing the probability for the variant carrying extra chromosomes to reach fixation is equivalent to computing the probability for the time difference to be negative (*t*_diff_ *<* 0). However, since the selection coefficient of aneuploids carrying extra chromosomes is lower than that of the euploid mutant, individuals of the former class will interfere with the expected progression of the aneuploid mutation to the fixation (see Fig. 1B), by an effect known as “clonal interference” (CI) [37–40].

We find that, to compute the probability to fix extra chromosomes, CI effects are captured by the extended condition 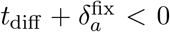, where 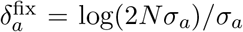 is the effective time to fixation of an aneuploid mutant carrying extra chromosomes [40] (i.e., the time interval during which CI effects can take place, see Material and Methods). This leads to the expression

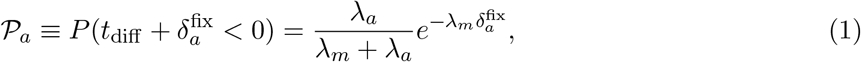

for the fixation probability. This expression is similar to the ones presented in [35, 41]. Consistently with ref. [37], CI effects are related to the expected number of euploid mutations that can emerge during the fixation dynamics of the mutant with extra chromosomes. In the limit 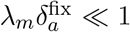, there are no interfering mutations and the probability to develop extra chromosomes is set by the fixation rates alone 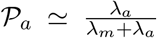 (Fig. 1C). In the clonal interference regime 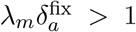 the emergence of aneuploidy with extra chromosomes is exponentially suppressed to zero (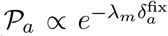, Fig.1D). Moreover, in this regime, the evolutionary dynamics would be characterized by the elimination of aneuploidy, resulting from the emergence of euploid mutants, after an initial increase in frequency of anuploid mutants. However, the observed loss of anueploidy would not signal the existence of karyotpe instability, as the duplicated chromosome would not be lost within aneuploid individuals. Our model can be exploited to investigate this scenario, and predicts the existence of a critical population size around which such dynamics (rise of aneuploidy to high frequency and subsequent elimination because of CI effects) could be observed (Fig. S9 and S.I.).

When 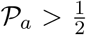, the emergence of aneuploidy is more likely than that of the competing beneficial point mutations. Hence, the condition 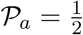 sets the a lower critical value for the beneficial selection coefficient

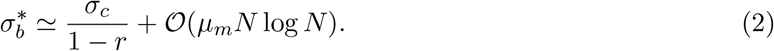

The prevalence of the evolutionary route developing extra chromosomes is observed in “stress” conditions where the beneficial effect exceeds the minimal value 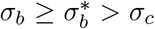. Here *r* = *μ*_*m*_*/μ*_*a*_ *<* 1 is the ratio between the mutation *μ*_*m*_ and the missegregation rate *μ*_*a*_ (see SI appendix).

### Aneuploidy with extra chromosomes is a “quick fix” in stressful conditions

The dynamics leading to the fixation of one of the two evolutionary routes can also be described in terms of the waiting time before the emergence of the fastest successful mutant. This dynamical quantity is described by the minimum of the two waiting times, *t*_min_ ≡ min(*t*_*a*_, *t*_*m*_), and has expected value (Fig.1E, see SI appendix for derivation)

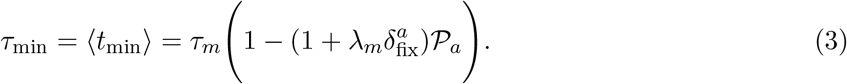

Thanks to the possibility of developing extra chromosomes, the waiting time until the emergence of the successful mutant is therefore shorter than the time needed to develop the competing set of point mutations *τ*_*m*_ = 1*/λ*_*m*_, which, in our model, would be attained if the mutational route was the only genomic change offering a solution to the external stress. This evolutionary route is still dynamically selected when *σ*_*b*_ *< σ*_*c*_ → *τ*_min_ *τ*_*m*_ = 1*/λ*_*m*_, i.e., when the global effect of extra chromosomes (benefit minus cost) is detrimental. Conversely, in the opposite limit *σ*_*b*_ *≫ σ*_*c*_ → *τ*_min_ *τ*_*a*_ = 1*/λ*_*a*_, the waiting time is set up by the fixation rate of extra chromosomes alone. Clonal interference effects 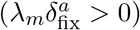 lead to an increase of the waiting time, i.e., reducing the speed of adaptation in response to the stress, consistently with refs [37–40].

In summary, the model describes in quantitative terms the early-stage evolutionary role of aneuploidy carrying extra chromosomes. According to the predictions, extra chromosomes provide a “quick fix” to the external stress (because 𝒫_*a*_ 1 → *τ*_min_ *τ*_*a*_ *< τ*_*m*_). Anaueploidy also has an indirect effect on the mutational dynamics of euploid individuals, by effectively selecting the fast mutants, hence causing a reduction of the waiting time to the emergence of the successful euploid mutant (1 *> 𝒫*_*a*_ *>* 0 → *τ*_min_ *< τ*_*m*_).

### The model correctly predicts the outcome of experimental evolution data from [20]

Our model can be applied to describe the evolutionary dynamics observed in experimental setups akin to ref. [20]. In their experiment, Yona and coworkers exposed four independent yeast populations of diploid strains to a constant heat stress of 39°C. After ∼ 450 generations, the duplication of chromosome III (trisomy) was found to have reached fixation in all four populations. The duplication of this chromosome was shown to carry a beneficial effect in response to the applied heat stress and to be the dominant evolutionary solution over an alternative mutational route attained by point mutations inducing the up-regulation of heat-shock genes.

In order to compare the model prediction to the outcome of the experiment, we obtained growth curves from the authors of [20], evaluated for the diploid and aneuploid strain (carrying the trisomy of chromosome III) both in normal conditions (30°C) and in stress conditions (39 °C). We used the growth curves to infer values of the selection coefficients of aneuploid individuals (see Materials and Methods, Fig.S1 and Table S1, the numerical values we obtained are *σ*_*b*_ = 0.17 gen^−1^ and *σ*_*c*_ = 0.05 gen^−1^).

Given these values for the selection coefficients, we evaluated the cumulative probability of developing aneuploidy (Eq. 13) *vs* time, according to our model prediction (Eq.s (1,3, see Material and Methods), using an effective population size of *N* = 10^6^ individuals, as a function of the missegregation rate (*μ*_*a*_) and of the total mutation rate (*μ*_*m*_) (Fig. 2). We find the model predictions Eq.s (1,3) to be in quantitative agreement with the outcome of the experiment, for realistic values of the missegregation rate (*μ*_*a*_ ≥ 8 * 10^−7^gen^−1^) and of the total mutation rate (*μ*_*m*_ ≤ 5 * 10^−9^). Similar results are obtained setting a bigger value for the effective population size of (*N* = 10, see Fig.S3).

**Figure 2:**
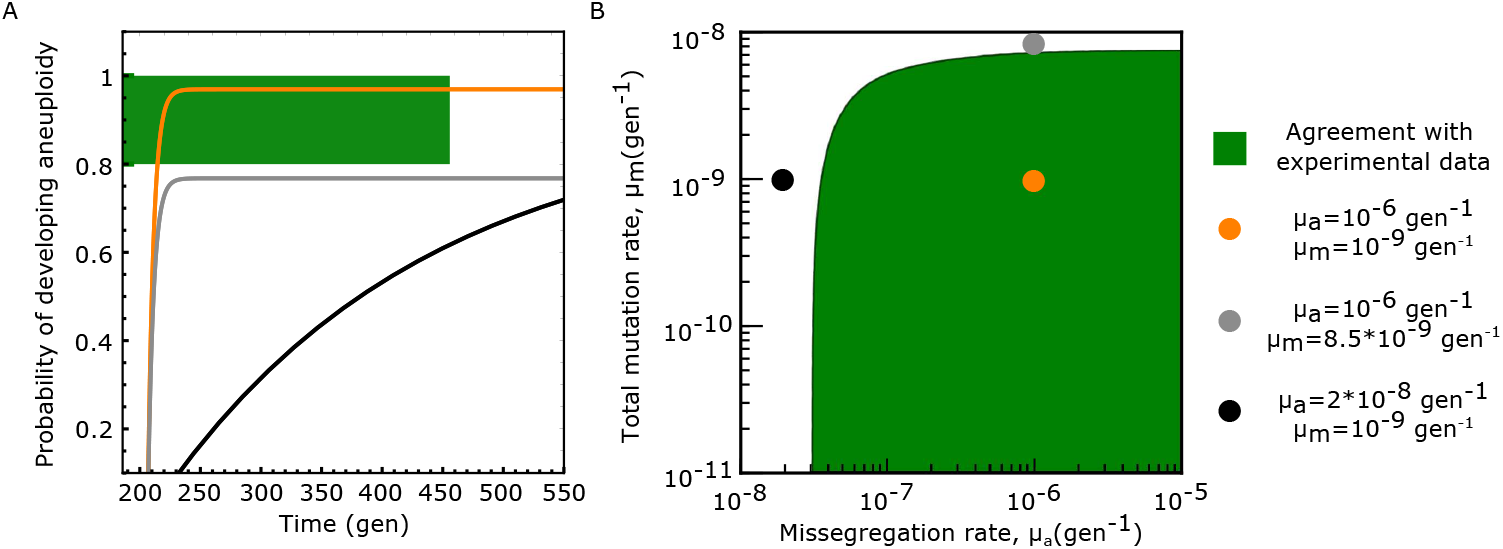
Model predictions agree with laboratory-evolution data from ref. [20]. (A) Expected cumulative probability for the emergence of aneuploidy with extra chromosomes *vs* the time to reach fixation (see Material and Methods), computed according to the model prediction (Eqs 1, 3) shown for three combinations of the values of the model parameters (*μ*_*a*_, *μ*_*m*_) (color coded, numerical values reported in the legend of the plot). In the experiment, where a yeast population was exposed to stress by increasing the temperature to 39*C*, 4 out of 4 yeast populations developed chromosomal duplications (*CI*_66%_ = [0.8, 1] for the probability to develop aneuploidy), and all the fixations were reached before 450 generations. Hence, the experimental data fall in region of the plot corresponding to *P*_*a*_ ∈ [0.8, 1] and *t* = 450*gen*, marked by a green bar and highlighted by green dashed lines. The trajectories predicted by the model that cross this region are in agreement with the experimental data. Similarly, panel (B) shows the combinations of the numerical values of the model parameters (*μ*_*a*_, *μ*_*m*_) that are in agreement with the experimental data. The colored circles mark the values of the model parameters that were used to generate the trajectories shown in A (each dot has the same color of the corresponding trajectory in A). Numerical values of the beneficial selection coefficient (*σ*_*b*_ = 0.17 gen^−1^) and for the fitness cost of aneuploidy (*σ*_*c*_ = 0.05 gen^−1^) were obtained from exponential fits of the growth curves of the corresponding yeast strains [20], (see Material and Methods and Fig. S1). The effective population size was set to *N* = 10^6^ individuals (Fig. S3 shows results for *N* = 10^7^). The data reported here refer to the “High-Temperature” experimental setup. Similar agreement between model prediction and experimental data is observed for the “High-PH” experimental setup (Fig.s S2)

In order to determine realistic ranges of the rates (*μ*_*a*_, *μ*_*m*_) in yeast, we reasoned as follows. The numerical value of the yeast per-base spontaneous mutation rate is *μ*_spont._ = 1.7 * 10^−10^gen^−1^ [42]. The mutation rate can be higher than spontaneous rate since the same phenotypic effect, i.e., the development of resistance to heat by up-regulation of heat-shock genes, can be attained with more than a single point mutation. A conservative estimate the size of this mutational target is no more than 100 bases, giving an upper bound constraint for *μ*_*m*_ ≤ 10^−8^gen^−1^. Values of the mutation rate lower than the spontaneous rate, on the other hand, would correspond to a scenario where the selection the euploid mutant does not develop a whole duplication of the expression of the target gene (but would alter gene expression to a lower degrees, see S.I.), and hence were also considered to be realistic. Measurements for the missegregation rate exist in the literature (*μ*_*a*_ 10^−6^gen^−1^).

The agreement between model prediction for the probability to develop aneuploidy and for the time fo fixation) and experimental data is observed yet another independent evolutionary experiment, described in ref.s [20,43], where a diploid yeast population was exposed to a different stressing environment, high PH. This experiment revealed the fixation of strains with the duplication of chromosome V (trisomy, see Supporting Information and Fig.s S1, S2, S3 and Table S1).

### The cost of extra chromosomes increases linearly with the total number of genes they contain

The fitness cost of an aneuploid strain is defined as the reduction of its per-individual offspring. In proliferative conditions, this can be proxied by the growth rate difference with respect to the euploid strain, evaluated in the same environmental conditions. An alternative proxy for fitness is the stationary-phase population size in a given condition. We deduced these proxies from both growth rates [9] and full growth curves [19] collected for yeast aneuploid strains grown in rich media and in the absence of external stress (see Material and Methods).

In both data sets, we found a statistically significant negative linear correlation between the growth rate of aneuploid strains and the total number of genes carried in the exceeding chromosome. This relation is observed (with different slopes) both in strains with disomic chromosomes compared to a haploid (ploidy 1) genomic background (Fig.3 A,B,C) and in strains with trisomic chromosomes compared to a diploid genetic background (Fig. 3 C). Notably the same trend is not only observed in aneuploid strains carrying only a single duplicated chromosome (Fig. 3 A), but also in strains with up to 8 duplicated chromosomes (Fig.3 B-C), suggesting that epistatic interactions between the fitness costs of multiple duplicated chromosomes are small. The data set from ref. [19] also shows a negative correlation between the fitness proxied by stationary-phase population size (optical density, OD) and the total number of genes carried in the excess chromosomes (Fig.3B). Linear negative correlations between growth rates and number of genes in excess chromosomes of aneuploid strains are also coherently observed in all the stress conditions investigated in the data-set from ref. [9] (Fig. S5 A).

**Figure 3:**
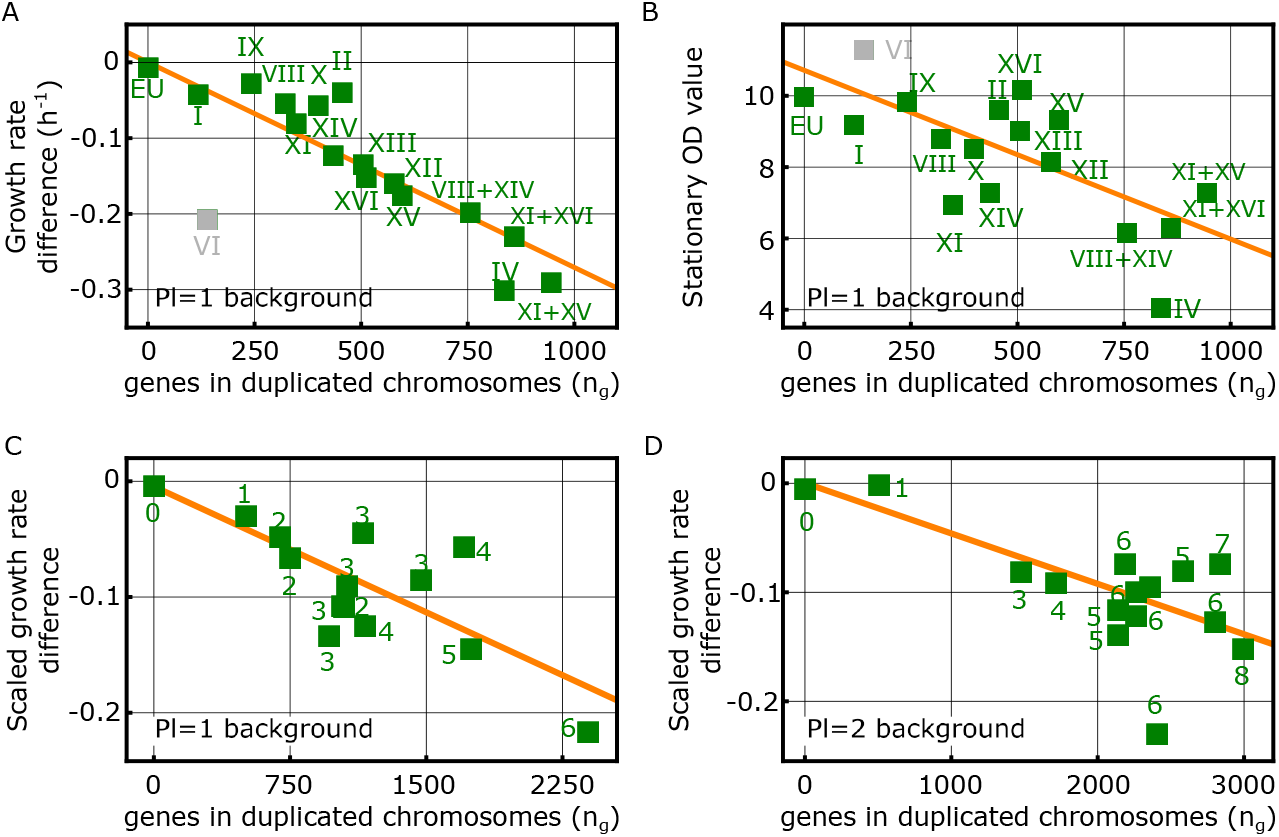
The fitness cost of extra chromosomes is proportional to the total number of genes present in the excess chromosomes. A: Plot of the values of the difference between the exponential growth rates of aneuploid strains with extra chromosomes and the exponential growth rate of the euploid strain (squares, labels indicating disomic chromosome numbers) against the number of genes carried in the disomic chromosomes (data from ref. [19], see Materials and Methods for details). The growth rate differences (estimating fitness differences) display a significant negative linear correlation with the number of duplicated genes (red line, Pearsons’ r = -0.93, p-value*<* 10^−6^). B: Values of the stationary-phase optical density (OD, squares, labels indicate disomic chromosomes) shown against the number of genes in the disomic chromosomes (data from ref. [19]). The stationary-phase OD of aneuploid strains, a complementary proxy for the fitness, displays a significant negative linear correlation with the number of duplicated genes (red line, Pearson’s r = -0.68, p-value*<* 0.005). In panels A and B, the data corresponding to the disomy of Chr VI was not included in the statistical evaluation, as this disomy in a euploid background is known to be lethal on its own [19, 21, 44, 45]. C and D: Scaled growth rate differences of aneuploid strains obtained from ref. [9] (see Material and Methods). The plots show scaled growth rates differences (squares) between aneuploid and haploid (C) or diploid strains (D) against the number of genes carried in unbalanced chromosomes (disomic chromosomes for the left plot and trisomic chromosomes in the right plot). Numbers next to the squares indicate the number of unbalanced chromosome carried in each strain. In both panels the proxied fitness difference of aneuploid strains display a significant negative linear correlation with the number of genes carried in extra chromosomes (red lines, Pearson’s r = -0.77 (B), -0.69 (C) and p-value≤ 0.001 (B), 0.004 (C).

Altogether, this experimental body of evidence suggests a general fitness cost for aneuploid individuals with extra chromosomes with respect to a euploid background, of the form

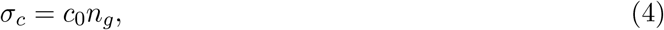

where *n*_*g*_ is the total number of genes carried in the extra chromosomes and *c*_0_ is the average cost per gene, which depends on the external condition and on the background. The fitness described by Eq. (4) does not necessarily imply that all the exceeding genes contribute to fitness, but also supports a scenario where only a fraction of the genes of the duplicated chromosome can reduce the reproductive fitness [46, 47]. The statistically significant linear correlations observed in Fig. 3, however, suggest that the subset of genes that contribute to the cost should be (roughly) evenly distributed across the genome. Indeed, only in such case the probability of finding a gene contributing to the fitness cost in a given chromosome would be proportional to its length, giving rise to the observed correlations. We note that this model considers chromosome copy number and duplication of different chromosomes equivalent in terms of per-gene cost. Specifically, the average fitness costs (*c*_0_) of aneuploid strains with diploid *vs* haploid background display a linear correlation (Fig. S5, B), suggesting the existence of a condition-specific effect on the fitness cost. Values of the fitness cost in the diploid background are found to be about a factor one-half of those observed in the haploid background, indicating that the development of extra chromosomes is suppressed in haploids and is more likely in diploids, an effect that is in agreement with observations based on evolutionary genomics data [21, 48, 49].

Of note, in Fig. 3A, the disomy of Chr VI shows the largest deviation from the linear decreasing trend (similar deviation was observed in [21]). This deviation results from an additional fitness cost that is specific of this disomy, which was reported to be lethal in the ploidy=1 background [19]. This additional cost is due to the two key cytoskeleton genes TUB2 (tubulin) and ACT1 (actin), which reduce cell viability when their expression is increased [44, 45]. Notably, the effect of the disomy of Chr VI is alleviated in combination with other aneuploidies, for example, Chr I and Chr XIII [19].

### A minimal fitness model for aneuploid strains with extra chromosomes

The analyses reported in Fig. 2 and 3 suggest that a global effect of extra chromosomes on the growth rate of a strain is recapitulated by a minimal fitness-landscape model, with no epistatic interactions between genes of the extra chromosomes. In this model, fitness is the sum of two contributions: (i) a fitness cost (*σ*_*c*_) that captures the empirical observation described by Eq.4 and (ii) a chromosome-specific fitness component, which captures the additional beneficial or detrimental effect of excess chromosomes *σ*_kar,*s*_. Under these assumptions, the selection coefficient of an aneuploid strain (*s*) in any given growth condition (environment or stress), with respect to the closest euploid background (haploid or diploid) takes the form

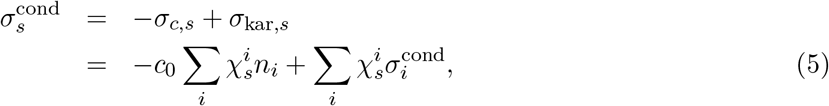

where the karyotype of the strain is defined by the characteristic matrix *χ*_**s**_, where 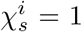 if, in the strain *s*, the *i*^*th*^ chromosome exceeds the background ploidy number. The fitness cost of the strain is due to the total number of exceeding chromosomes, 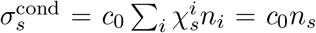, where *c*_0_ *>* 0 is the condition-specific average fitness cost per gene, *n*_*i*_ is the number of genes in the *i*^*th*^ chromosome and *n*_*s*_ is the total number of extra chromosome of strain *s*. Each aneuploid chromosome has an effect on the growth rate 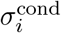, which can either be beneficial 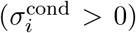 or detrimental 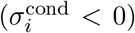 and is condition-specific. This results in the kariotype fitness component 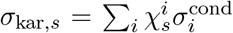. A condition (environment or stress) is defined by the value *c*_0_ and the set of values 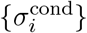.

### The fitness landscape defined by Eq. 5 captures nontrivial behavior of stress phenotypes

The two fitness components of the minimal model, i.e., the fitness costs *σ*_*c,s*_ and the chromosome specific fitness effects *σ*_kar,*s*_, can be inferred from large-scale studies of anuploid yeast phenotypes in stressful conditions, such as ref. [9] (see Material and Methods).

The first component captures the global linear decreasing trend of the growth rates of aneuploid strains *vs* the total number of exceeding genes, as discussed above. Interestingly, this component can also explain in quantitative terms the observed linear correlation between the degree of phenotypic variation and the degree of overall growth suppression, observed in the data [9] and modelled in refs [28, 30]. In our modelling framework, this correlation corresponds to a linear relationship between the average value 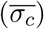 and the standard deviation 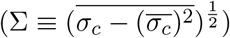 of the fitness cost evaluated in a cohort of aneuploid strains. Here, we have denoted with 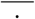. averages computed over the cohort of aneuploid strains in a given growth condition. The fitness cost Eq. 4 predicts a linear relationship of the form

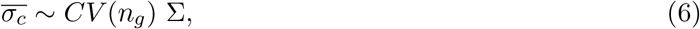

where 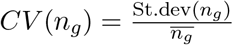 is the coefficient of variation of the distribution of the number of exceeding chromosomes (the total number of genes contained in aneuploid chromosomes) evaluated in the set of aneuploid strains considered. The quantitative expression Eq. 6 explains about 80% of the observed variability of the growth rates in the data-set of ref. [9], implying that the fitness cost alone cannot explain the whole range of observed phenotypic diversity (Fig. S6).

The deviations from this linear trend are captured by the second (condition- and chromosome-specific) fitness component (*σ*_kar,*s*_), where the effect of an aneuploid extra chromosome (*i*) on the growth rate is quantified by a chromosome- and condition-specific fitness effect, 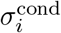. Fig. S7A reports the inferred values of the of the fitness-gain component of each chromosome across stress and control growth conditions for the Pavelka *et al*. data set. These inferred values are net of possible confounding factors due to the per-gene fitness cost highlighted previously. Curiously, the chromosome-specific fitness components in the same environment are generally different (uncorrelated) between the ploidy=1 and ploidy=2 backgrounds (Fig. S7B). This difference could suggest that the effect of chromosome duplication is ploidy-specific, consistently with observations of other ploidy-specific fitness effects [50]. However, since in the Pavelka *et al*. experiments each strain was carrying more than a single aneuploidy, suggesting the presence of epistatic effects between different added chromosomes.

Unfortunately, the current data are too sparse to infer such epistatic interactions. Additionally, the difference of the chromosome-specific fitness effect between the ploidy=1 and ploidy=2 background could be related to different physiological constraints seen by haploids and diploids, and to the different relative gene-dosage increase resulting after a duplication in the two different backgrounds.

Looking at Fig. S7A, one clearly sees that adding different specific extra chromosomes can improve or decrease the fitness in a specific environment, but each environment is characterized by the extent of such fitness gains and losses. For example, in stressful environments such 4NQO, adding an extra chromosome to the genetic background could improve or decrease the fitness by a factor that is more than 10-fold larger than performing the same operation in a non-stressful condition such as glycerol media. Because of this property, it is tempting to classify the “harshness” of an environment by the variability in behavior of aneuploids bearing specific extra chromosomes. Indeed, the variability of effects across chromosomes in a fixed given condition is found to be proportional to the fitness cost per gene observed in the same environment (cfr. Fig. S8ABCD), which can be seen as an independent evaluation of the harshness of that environment. Additionally, contrary to the effects of specific chromosomes, the distributions of the fitness components for the ploidy=1 and ploidy=2 backgrounds (shown in Fig. S8CDE) share common properties that are related to the growth condition. In particular, each environment is characterized by distributions of chromosome-specific fitness effects that have a similar width for ploidy=1 and ploidy=2 backgrounds (Fig.S8E).

### Expected inter-population dynamics of aneuploids

The minimal fitness-landscape model described by Eq.5 can be used to describe the expected inter-population dynamics of aneuploid strains with a single chromosome gain, by investigating the substitution dynamics associated to Eq. 1 in the landscape (Eq.5) when 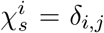, for some *j >* 0, and *δ*_*i,j*_ is the standard Kronecker delta. Following a standard population dynamics approach [51, 52], we can use a probabilistic framework to characterize the selective effects of a generic environment on the growth rate of a strain with an aneuploid chromosome, by assuming that the beneficial effect (*σ*_*b*_) is exponentially distributed, 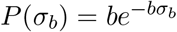. Averages with respect to this distribution, denoted with ⟨⟨. ⟩⟩, quantify the expected dynamics of aneuploid strains with excess chromosomes in a set of conditions. Hence, they can be used to generate predictions on the typical population dynamics of aneuploids. Under these assumptions, the model predicts that the average probability of developing aneuploidy with extra chromosomes,

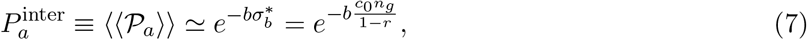

decreases exponentially with the number of genes contained in the extra chromosomes, suggesting in particular that the relative abundance of duplicated chromosomes is exponentially suppressed with their length. In addition, the typical selection coefficient of an aneuploid strain that has reached fixation,

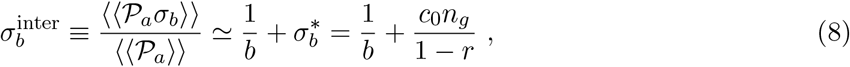

is expected to increase linearly with the cost of extra chromosomes, implying in particular that longer chromosomes require higher fitness advantage to reach fixation.

Combined together, the two model predictions (Eq.s 7 and 8), suggest an “equilibrium” distribution of aneuploid strains of the form (see Material and Methods)

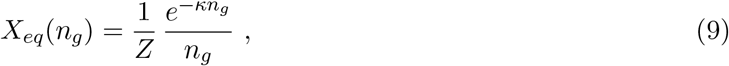

where *n*_*g*_ is the number of genes contained in the aneuploid chromosome, *Z* is a normalization factor, and *κ* is an effective fitness cost per gene (see Material and Methods). This prediction is in good agreement with the relative abundances of yeast aneuploid strains observed in evolutionary genomics data (Fig. 4). Interestingly, the numerical values of the effective fitness cost per gene are in agreement with existing experimental evidence [53, 54] suggesting a reduced fitness cost for wild strains (collected as “natural strains” in refs. [53, 55] as “wild strains” in ref. [54]). In other words, these strains have a higher propensity to generate aneuploidy, when compared to strains of other kinds, including domesticated, industrial and human-associated strains (cfr. Table S3). We find similar results when comparing the abundance of strains with a ploidy¿2 background to that of strains with a lower ploidy background, finding that extra chromosomes are associated to a lower fitness cost in a ploidy¿2 background (cfr. Table S3).

**Figure 4:**
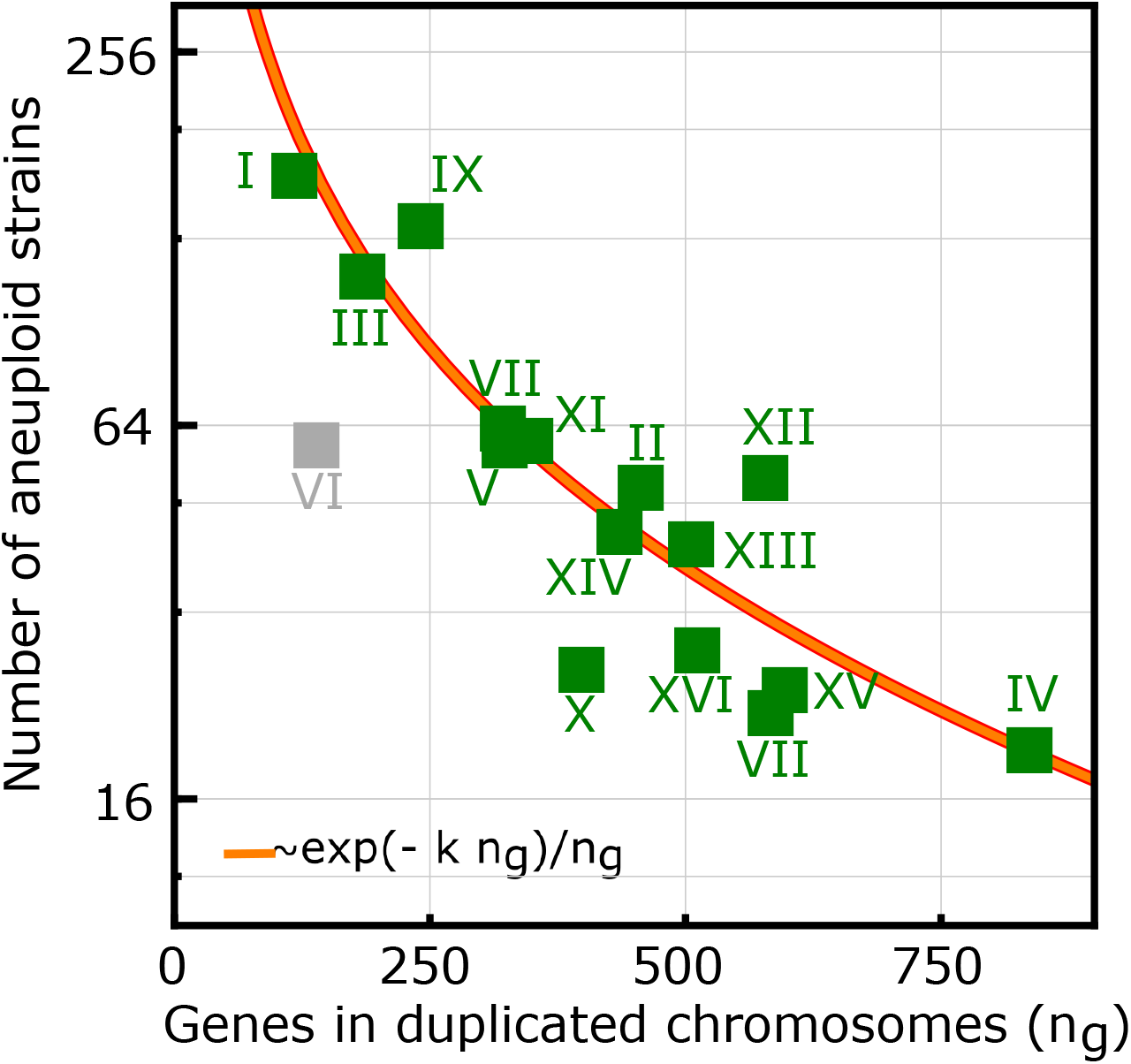
The fitness landscape derived from phenotypying of laboratory yeast strains explains the relative abundances of yeast aneuploid strains observed in evolutionary genomics data. Relative abundances of aneuploid strains vs the number of excess genes contained in the aneuploid chromosome (squares). The numbers of aneuploid strains were retrieved from published data collected in eight studies and reported in ref. [21](cfr. Table S3). The orange line shows the fitness model expectation, for the relative equilibrium frequencies set by chromosome acquisition and loss rates of Eq.s (7,8), which predicts a functional dependence of the relative frequencies on the number of excess genes (*n*_*g*_) of the form ∝ exp(−*κn*_*g*_)*/n*_*g*_. Numerical values of model parameters are reported in Table S3. Data-count of the duplications of Chr VI (gray square) was not considered in the model fit, since this chromosome is known to be lethal because of the specific effects of the main cytoskeletal genes tubulin and actin [19, 21, 44, 45].

## Discussion

In yeast, the development of aneuploidy resulting from an accidental chromosome missegregation has been characterized with massive experimental data [3–6, 6–10, 19, 20, 20–22]. As a consequence of this major effort, we are in need of unifying principles to rationalize this wealth of data, and embed the underlying evolutionary dynamics into simple quantitative models. Here we have focused on a specific question, the role of chromosomal duplication with respect to a reference euploid background. Our results show that a simple evolutionary model where a fitness cost of chromosome duplications is counterbalanced by a fitness advantage from the expression of specific genes can explain in quantitative terms two key observations of the emergence dynamics of aneuploidy: (i) chromosome duplications emerge transiently as a “quick fix” to dosage insufficiency of a single gene in stressful environments [11, 19, 56] (ii) depending on the nature of the applied stress, aneuploidy or local mutations may be favored [20].

While traditionally the fitness advantage of a phenotype associated to a certain mutational target was considered to be the primary trait related to its adaptive value, the recent debate has challenged this assumption based on experimental results that highlight an important role of mutational paths and mutation rates. Our analysis of aneuploids with extra chromosomes provides another example where mutational paths with high rates may give a more relevant contribution to adaptation than mutations with large benefits occurring more rarely [33–35]. Our results support the existence of a cost of single-chromosome duplications that is proportional to the number of genes contained in the exceeding chromosomes. This simple behavior is surprising, due to the numerous documented complex physiological changes that emerge with aneuploidies, such as dosage imbalance, effects on interaction networks, and consequent osmotic effects [7, 57]. Importantly, our results are in line with a scenario where only a fraction of the genes in a duplicated chromosome will actually contribute to the fitness cost, as supported by the results of refs. [46, 47]. These studies also find that the fitness costs can be complex and specific of a genetic background, and that in many cases they are due to stoichiometric imbalance between proteins that are interaction partners. More specifically, this scenario implies that some duplicated genes give a null contribution to the fitness cost, hence, the average fitness (*c*_0_) should be thought as the average of a bimodal distribution (genes with a non-null contribution plus genes giving a null contribution). Moreover, our analysis (Fig. 3) would suggest the class of genes contributing to the cost to be evenly distributed across the genome. This model interpretation is in agreement with existing literature [46, 47, 58–60]. Moreover, the remaining fraction of genes that do not contribute to the cost, would be described by our model as behaving neutrally, and could therefore be retained in small segmental duplications; this aspect would be in agreement with the conclusions of ref. [58].

Another interesting interpretation of this form of the fitness cost is that the reduction of the growth rate of aneuploid strains may be the result of the interdependence between growth rate and gene expression, in accordance to the one described by phenomenological laws first observed in bacteria [61], and more recently also in yeast [62]. A possible role of resource allocation in the fitness cost of overexpressed proteins was also suggested in a direct investigation of the cost of overexpressed proteins [47], which, however also found that these effects vary considerably with the genetic background. Genes contained in an extra chromosome are unnecessary for the survival and their expression induce a reduction of the growth rate by effectively decreasing the fraction of resources allocated to the ribosomal and housekeeping protein sectors, leading to a decrease in growth rate. This connection holds only if the genetic gene dosage is proportional to gene expression, an effect experimentally observed in for yeast [9, 19, 63]. Interestingly, the connection between the fitness cost and gene dosage is coherent with our analysis, since we observe that values of the fitness cost in the diploid background are close to one-half than of those observed in the haploid background, suggesting a connection of the fitness cost per gene to the relative extra gene dosage. Importantly, the linear (per-gene) cost of duplicated chromosomes appears to be a common unifying feature of the fitness landscape in different conditions and environments.

Our model assumes that, when duplicated, some genes will impose a fitness cost to the cell, causing a reduction of its proliferation rate. However, the biological mechanisms that cause this cost are not described within the model, leaving room for several interpretations and further modeling efforts. One contribution to this cost is related to regulatory effects and could be associated to all the genes that are up-regulated together with a duplicated gene, as a result of both cis- or trans-regulations, and additionally, as discussed above, stoichiometric imbalances in protein-interaction networks caused by dosage changes are found to play an important role (by regulatory as well as biophysical effects). For this contribution, the proportionality observed in (Fig.3) would suggest the number of genes affected to be proportional to the number of duplicated and costly genes (i.e., the number of genes that are duplicated and contribute to the cost), meaning that what matters for the main trend is the average number of interaction partners of each such gene.

The formulation of a minimal fitness-landscape model (Eq.5) informed by data allows for the inference of chromosome-specific fitness effects. Such effects are only related to the dosage increase of genes contained in specific duplicated chromosomes, offering a quantiative framework for the inference of fitness components of early aneuploids. In addition, we have shown that a simple description of the longer-term evolutionary dynamics of our model (Eq.s1,3) in this landscape captures the relative abundance of aneuploidies observed in yeast population genomics data. Hence, this model can be used to investigate both the intra-population dynamics of aneuploidy individuals within an evolving population (cfr. Fig.2) and the substitution dynamics at the inter-population population level (cfr. Fig.4).

Importantly, our model can be used to design evolutionary experiments to investigate key biological questions related to the emergence of aneuploidy, which require precise quantitative assessments. For example, our model could be used to test whether the karyotype state of aneuploid individuals is stable in the long term (and in which conditions). As we have shown (see Fig.S9 and SI Appendix), for this question to be address it would be important to design experiments where Clonal Interference effects would be expected, since an observed dynamics with initial rise of aneuploids followed by its elimination from the population could be misinterpreted as a signal of kariotype instability, while simply being be the signature of CI. Moreover, our modeling framework could be deployed to design and investigate Mutation Accumulation (MA) experiments aimed at measuring the mis-segragation rate. In particular, our quantitative expression for the fitness cost of aneuploid individuals (see Fig.3 and and Eq.4) could be use to account for fitness effects in MA setups and correct the estimate of the missegregation rate [64, 65].

The model introduced here can be extended to describe more complex scenarios. First, it can be applied to investigate the evolutionary consequences of a sudden increase of the mis-segregation rate [6], which can result from the usage of anti-mitotic drugs. Second, it can be used as an ingredient to build models of chromosomal instability, with a clear interest for cancer development [66]. For the latter aspect, it would be important to clarify to what extent the increase of gene dosage translates into protein production in human cells [22].

## Methods

### Simulations of the evolutionary model

We performed numerical simulations of a standard Wright-Fisher model with mutations and selection, with constant population size *N*. Individuals of the populations are grouped into three distinct and non-overlapping classes: (a) euploid individuals, (b) aneuploid individuals and (c) euploid individuals with point mutations. Class (b) is generated from class (a) with a rate *μ*_*a*_ (per individual, per generation) and its members have a selection coefficient *σ*_*b*_ −*σ*_*c*_. Similarly, individuals of class (c), characterized by a selection coefficient *σ*_*b*_, are generated from individuals of class (a) with a rate *μ*_*m*_ (per individual, per generation). The simulation is initialized with all individuals assigned to class (a) and it is stopped when either class (b) or class (c) reaches a frequency *x* ≥ 0.95. At the end of each simulation, we recorded the successful class (either (b) or (c)) and the time of appearance (measured in generations) of the first mutant whose descendants took over the whole population (i.e., the emergence time of the their last common ancestor *t*_min_).

### Evolutionary parameters for the experimental data

**[20]**.To quantify the fitness cost of the aneuploid strain investigated in [20], we made use of growth curves of the aneuploid and the euploid strains, evaluated at permissive conditions, i.e., without stress. We then inferred the growth rates 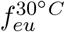 and 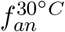 with an exponential fit of the corresponding growth curves (Fig. S1 A). Similarly, for evaluation of the fitness benefit of the anuploid strain in presence of a heat stress (39°C), we inferred the growth rates 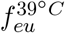 and 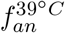 (Fig. S1 B). The two selection coefficients were then computed from the following set of equations

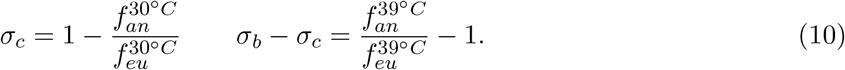

Selection coefficients of the aneuploid strain for the experiment performed in high pH were computed analogously, using growth curves of the aneuploid and euploid strain evaluated in permissive (Fig. S1 C) and stress condition (Fig. S1 D). Numerical values for the inferred growth rates are shown in Tab.S1.

To estimate the effective population size of the wells used during the evolution experiment, we used the following argument. The experiment in [20] was performed in 96 well plates, with a max volume per well ≃0.4*ml*. During the experiment, cell density in liquid cultures was monitored by optical density at 600 *nm*, reached a maximum value ∼ 1 *OD*_600_. Since the value *OD*_600_ = 1 corresponds to approximately 10^7^ cells per ml [67], we estimated the effective population size to be 10^7^ ⪆ *N ⪆* 10^6^. In Fig.2 and Fig.S2 we show results obtained with *N* = 10^6^ while in Fig. S3 we show equivalent results computed for *N* = 10^7^.

The expected cumulative probability for the emergence of aneuploidy with extra chromosomes was computed as using two model ingredient. First, the cumulative distribution describing the probability to have a successful aneuploid mutant (i.e., a mutant that eventually will reach fixation) emerging before time *t*, which reads

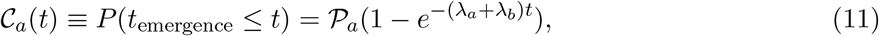

where 𝒫_*a*_ is the aneuploidy fixation probability (Eq.1), while *λ*_*m*_ = *μ*_*m*_*Nϕ*(*σ*_*b*_, *N*) and *λ*_*a*_ = *μ*_*a*_*Nϕ*(*σ*_*b*_ − *σ*_*c*_, *N*) are the fixation rates for the euploid and the aneuploid mutant. The fixation probabilities are computed according to Kimura’s expression *ϕ*(*σ, N*) = (1 − *e*^−2*σ*^)*/*(1 − *e*^−2*σN*^) [36]. Second, the time to fixation of the anuploid mutant, which reads (see SI and ref [40])

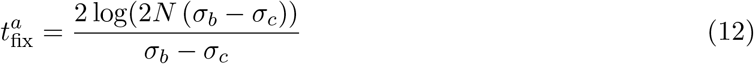

Finally, the expected cumulative probability for the emergence of aneuploidy reads

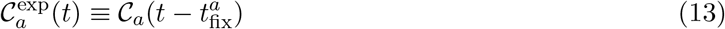

### Growth curves data from [19] and inference of growth rates

In the data-set collected by [19] yeast strains were grown in liquid cultures, and *OD*_600_ measurements were taken for several time points. Aneuploidy strains were engineered to harbour two specific genes (HIS3 and KAN), integrated in the two copies of the disomic chromosomes (one per copy). The two genes were also integrated in two chromosomes of the euploid strain. Growth curves were evaluated in media that is selective for the two genes (–His+G418 medium), therefore preventing the loss of one of the two disomic chromosome in the aneuploid strans, but is otherwise be neutral for other traits and does not induce a fitness difference between euploid and aneuploid strains.

Growth rates were then inferred fitting the growth curves to a logistic model

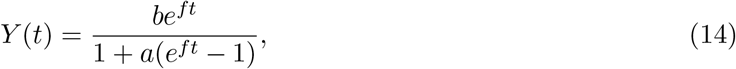

where *f* is the growth rate of the exponential phase, *b* sets the initial condition *Y* (0) = *b*, and 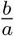 quantifies the fitness in the stationary phase of the growth (max OD value). We have fitted the data with a parametric Bayesian Model log(*Y* (*t*)) ∼ *𝒩* ((log(*b*)+*f t*−log(1+*a*(*e*^*ft*^ −1)), *σ*_*Y*_), choosing priors *a* ∼ *𝒰* (0, 1), *b* ∼ *𝒰* (0, 1), *f* ∼ *𝒰* (0.1, 1.1) and, 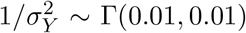, where ∼ stands for distributed as, and 𝒩, 𝒰, and Γ stand for normal, uniform, and gamma distribution. Model fits to the data are shown in Fig.S4 and inferred model parameters are summarized in Table S2.

### Data-set from ref. [9] and evaluation of growth rate differences

Yeast strains in the dataset collected by [9] were grown on solid media plates and growth data were obtained by automated spot detection and intensity measurements. The data-set consisted in 38 fully isogenic aneuploid yeast strains with distinct karyotypes and genome contents between 1*N* and 3*N*, and 3 strains euploid strains (one for each ploidy). In our analysis, we retained strains whose karyotype can be identified as an aneuploid resulting from chromosome gain, hence, we required the total number of gene contained in the excess chromosomes to be about one half of the total number (≤ 3200 genes). Aneuploidy strains of this form included the majority of the strains of the original dataset (14 strains with a ploidy=1 background and 15 strains with a ploidy=2 background). Since our analysis involved a stratification of the data according to the closest euploid strains, aneuploid strains with a ploidy=3 were discarded as too few for a statistical investigation (less than 5 strains).

Growth rates differences were evaluated as follow. The data-set consisted of values of the Optical Density (OD) of growth assays, evaluated at the same time (*t*_*max*_), of a set of strains with a similar initial number of cells (*N*_0_). Assuming exponential growth (growth assays that reached saturation were excluded from the analysis by the authors), the OD of a specific strain (*s*), at a given growth condition (*c*), can be modelled as

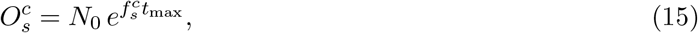

where 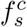 is the growth rate of strain *s* in the growth condition *c*. OD Values were then normalized to the value observed for the euploid strain with a ploidy=1 background, and in the same growing condition, obtaining transformed values

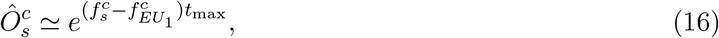

which we have log-transformed to get

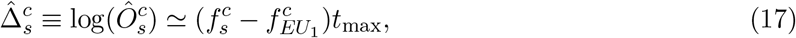

which are scaled (a-dimensional) growth rate differences between a given strain *s* and the euploid (pl=1) controll strain, evaluated in the growth condition *c*. Scaled growth differences w.r.t. the closest strains were then obtained by difference

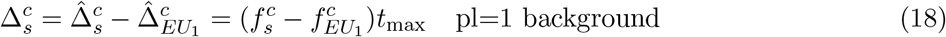

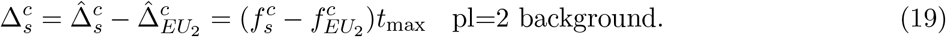

These values were used to evaluate statistics related to the fitness cost component shown in Fig.2 and Fig.s S5 andS6, as well as for the inference of the chromosome fitness components shown in S7 andS8.

### Inference of fitness components from data-set from ref. [9]

We consider a minimal fitness model, where the growth rate 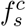 of an aneuploid strains *s* in the growth condition *c* reads

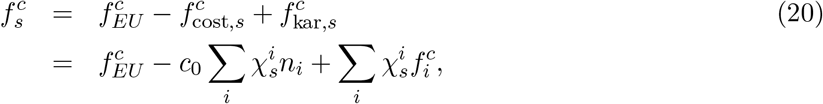

where 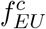 is the growth rate of the closest euploid strains to *s* in the same condition. The karyotype of the strain is defined by the matrix *χ*, where 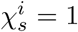 if, in the strain *s*, the *i*^*th*^ chromosome exceeds the background ploidy number. The fitness cost of the strain is due to the total number of exceeding chromosomes, 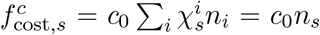, where *c*_0_ *>* 0 is the condition specific average fitness cost per gene, *n*_*i*_ is the number of genes in the *i*^*th*^ chromosome and *n*_*s*_ is the total number of exceeding chromosome of strain *s*. Each aneuploid chromosome has an effect on the growth rate 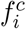, which can either be beneficial 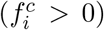 or detrimental 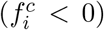 and is condition specific, that results in the kariotype fitness component 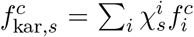. In the minimal model Eq.21 epistatic interactions between chromosomes are not considered.

For each growth condition of the data-set of Pavelka et al, we have inferred the model parameters *c*_0_ and the chromosome fitness effects 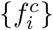 as follows. It should be noted that, while Eq.21 requires growth rates (in units [time]^−1^), the data-set by Pavelka et al consisted of scaled, a-dimensional values 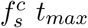, where *t*_*max*_ is a value that is constant for all the strains considered (the time duration of the growth assay). The model Eq.21 can therefore be inferred with the considered data-sets, since all the growth rates are scaled by the same value, and the inferred model parameters of Eq.21 will be expressed in a-dimensional units.

To estimate the value *c*_0_ we performed a linear fit of the data 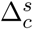 (Eq.19, Matherial and Methods) vs *n*_*s*_, inferring the linear model

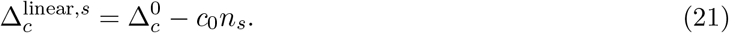

The value of 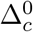 is a correction to the fitness of the euploid strain. Since the data was normalized to the growth of the euploid strain with pl=1, in the pl=1 data set we imposed 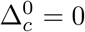 while in the pl=2 the parameter was set free.

Deviations of the data from the linear model were then used to infer the chromosome fitness effects. We first subtracted the linear model contribution, obtaining the detrended data

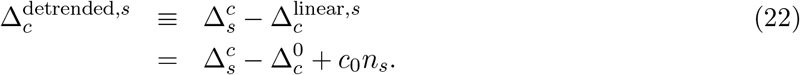

The chromosome fitness component are the solution of the linear system

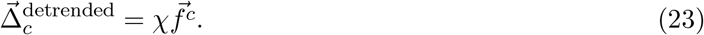

Since the matrix *χ* is sparse, the system of equations Eq.23 can not be solved exactly. Hence, we use the approximated Least Square solution of Eq.23

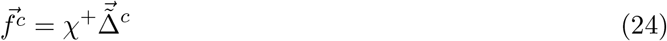

where *χ*^+^ is the pseudo-inverse of 𝒳, the matrix that specify the kariotypes of the strains considered.

Inferred values of the chromosome fitness components are shown in S7 and S8. Fig. S10 shows a detailed example of the inference procedure discussed in this paragraph.

### Fitness-landscape prediction for the relative abundances of aneuploid strains

We computed the equilibrium distribution for the relative abundances of aneuploidy strains as the ratio between the onset rate (*r*), i.e., at which aneuploid strains reach fixation, and the loss rate (*l*), i.e., the rate at which a aneuploidy is lost because euploid individuals reach fixation,

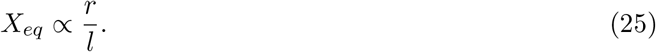

In this context, aneuploidy strains are identified by the number of genes that are contained in the duplicated chromosome, hence the only dependence on the chrosomome identity is via its gene content (*n*_*g*_). The two rates corresponding to the model considered here (see Fig.1) are defined in terms of the model predictions Eq.s (78) as follows.

The onset rate can be written as the product of 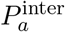, the intra population fixation probability (Eq.7), and an effective rate *μ*_stress_, describing the rate at which yeast population are exposed to stress conditions that can promote the emergence of aneuploidy

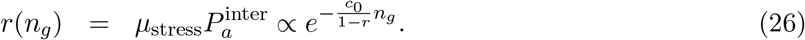

The loss rate rate depends on the environmental condition. If the stress condition that promoted the emergence of aneuploidy is no longer in action, then the population will restore the original euploid strain by losing the duplicated chromsome. In this case the euploid strain, generated with a missegragation rate per individual 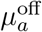, has a beneficial selection coefficient *σ*_*eu*_ = *σ*_*c*_(computed w.r.t. the anuploidy individual) and the loss rate will equal the substitution rate

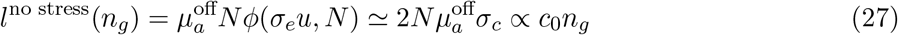

while *ϕ*(*σ, N*) = (1 − *e*^−2*σ*^)*/*(1 − *e*^−2*σN*^) is the Kimura’s fixation probability [36]. If the stress condition persists, then in the long term the population will substitute aneuploids with euploid with point mutations, i.e., the second mutational channel considered in our model. In the case the loss of aneuploidy would be attained by the sequential generation of an euploid individual, with a missegragation rate per individual 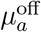 and selection coefficient 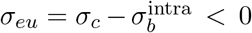, which then generates a mutant with a mutation rate individual *μ*_*m*_ and selection coefficient *σ*_*m*_ = *σ*_*c*_ *>* 0.Note that selection coefficients are now computed w.r.t. the anuploidy individual. These two sequential events are known to take place through the so called “stochastic tunneling” process [68, 69], that makes possible progression through intermediate deleterious alleles without the population ever experiencing the transient decline in fitness that would necessarily occur with sequential fixation. Hence, in presence of stress, the offset rate is equal to the tunneling rate [68–71]

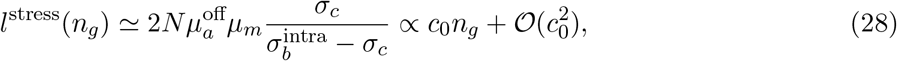

where we used the expression 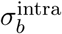 as in Eq.(8).

While the exact form of the equilibrium distribution will differ if considering persisting/non persisting stress conditions after the fixation of aneuploidy, it can be expressed in general terms as a scaling law (*vs* s the number of genes contained in the aneuploid chromosome *n*_*g*_) that is valid in both conditions, and takes the form

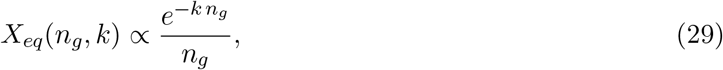

where 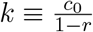 is defined in terms of the condition-specific fitness cost per gene (*c*_0_) and the mutational bias towards the generation of anuploidy individuals *r* = *μ*_*m*_*/μ*_*a*_ (see Main Text). To account for the variability of the growing conditions, which reflects in the variability of the parameter *k*, we assume the set of environmental stresses to be described by a uniform distribution in for *k* ∈ [0, 2*κ*], where 2*κ* is an upper bound for *k*. By averaging Eq. 29 over this distribution we find

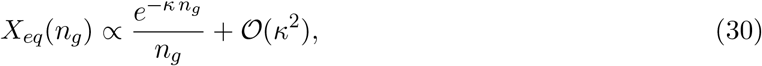

whose normalized form corresponds to Eq. 9. The value of the parameter *κ* is an effective fitness cost per gene, which is proportional to the max value of *c*_0_ of the set of growing conditions considered. We note that the numerical value of *c*_0_ is expected to be lower than the inverse of the typical chromosome size (the longest chromosome in yeast has *n*_*g*_ ≃ 600 genes), supporting the approximations taken in Eq.s (28, 29). This approximation is also validated a-posteriori, by the numerical values obtained in the model fit (cfr. Tab.S3).

## Aknowledgements

We would like to thank Andrea Ciliberto, Gilles Fischer, Gianni Liti, Bertrand Llorente, Rong Li and Paolo Bonaiuti for useful discussions. We are also very grateful to Avihu Yona, Eduardo Torres, Maitreya Dunham, Norman Pavelka and Giulia Rancati for having shared their experimental data with us. This work was supported by Associazione Italiana per la Ricerca sul Cancro AIRC IG grant no. 23258 (M.C.L. and SP). S.P. was supported by Fondazione Umberto Veronesi.

---

**Figure S1:**
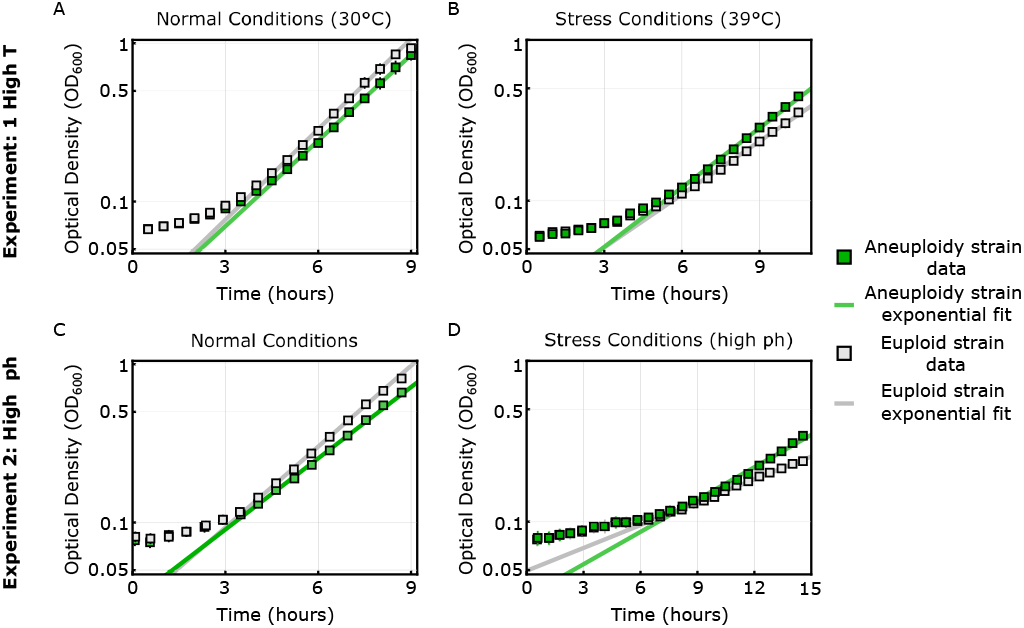
Growth rates describing of the aneuploid and euploid strain from ref. [20] are inferred from growth curves. Shown here are growth curves (Optical Density vs time) of the aneuploid strain (diploid strain with the trisomy of chromosome III) and of the diploid strain with the same genetic background evaluated both in normal conditions (30°C, A) and in stress conditions (39 °C, B). Similarly, in C and D we show the growth curves of the aneuploid strain aneuploid strain (diploid strain with the trisomy of chromosome IV) and of the diploid strain with the same genetic background evaluated both in normal conditions (C) and in stress conditions (high ph, D) The experimental data (squares + bars, showing mean and standard deviation evaluated over a set of ∼ 35 replicates) is shown together with an exponential fit (solid lines), evaluated neglecting the lag phase (∼ 4 hours in A, ∼ 5.5 hours in B, ∼ 4 hours in C and ∼ 7.5 hours in D). Values of the inferred growth rates are reported in Table S1.

**Figure S2:**
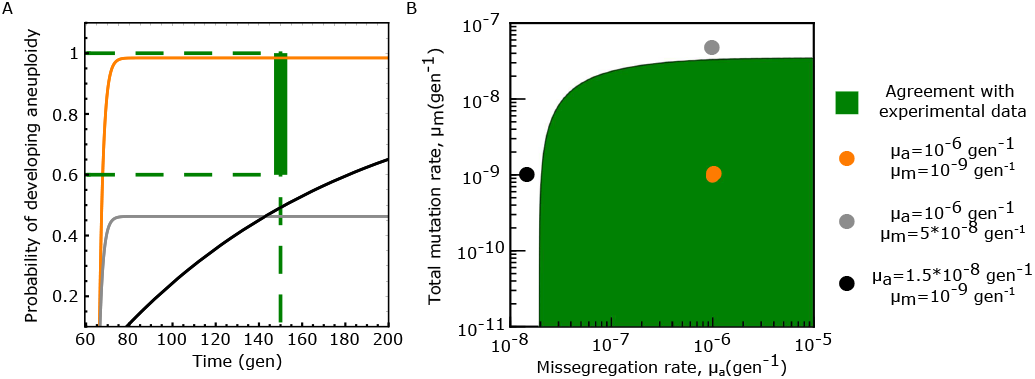
Model predictions agree with laboratory-evolution data from ref. [20] (High ph experimental setup). (A) Expected cumulative probability for the emergence of aneuploidy with extra chromosomes vs the time to reach fixation (see Material and Methods), computed according to the model prediction (Eqs 1, 3) shown for three combinations of the values of the model parameters (*μ*_*a*_, *μ*_*m*_) (color coded, numerical values reported in the legend of the plot). In the experiment, where a yeast population was exposed to stress by increasing the temperature to 39*C*, 1 out of 1 yeast population developed chromosomal duplications (*CI*_66%_ = [0.6, 1] for the probability to develop aneuploidy), and the fixation wes reached before 150 generations. Hence, the experimental data fall in region of the plot corresponding to *P*_*a*_ ∈ [0.6, 1] and *t* = 150*gen*, marked in green, delineate. Trajectories predicted by the model that cross this region are in agreement with the experimental data. Similarly, in (B) we show the combinations of the numerical values of the model parameters (*μ*_*a*_, *μ*_*m*_) that are in agreement with the experimental data, while coloured dots marks the values corresponding to the trajectories shown in A. Numerical values of the beneficial selection coefficient (*σ*_*b*_ = 0.29 gen^−1^) and for the fitness cost of aneuploidy (*σ*_*c*_ = 0.12 gen^−1^) were obtained from exponential fits of the growth curves of the corresponding yeast strains [20], (see Material and Methods and Fig. S1). The effective population size was set to *N* = 10^6^ individuals (Fig. S3 shows results for *N* = 10^7^).

**Figure S3:**
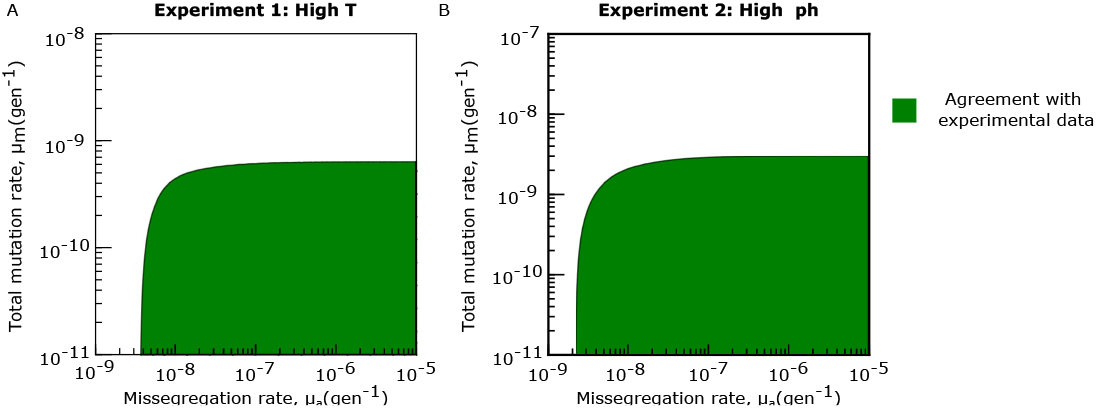
Quantitative model predictions agree with the laboratory evolutionary experiment from ref. [20] for *N* = 10^7^. Values of the model parameters (*μ*_*a*_, *μ*_*b*_) for which the model prediction is in agreement with the experimental data of ref [20] when assuming a population size of *N* = 10^7^ individuals, for the experimental setup at high temperature (A) and high ph(B) of agreement b as Fig.**??**B

**Figure S4:**
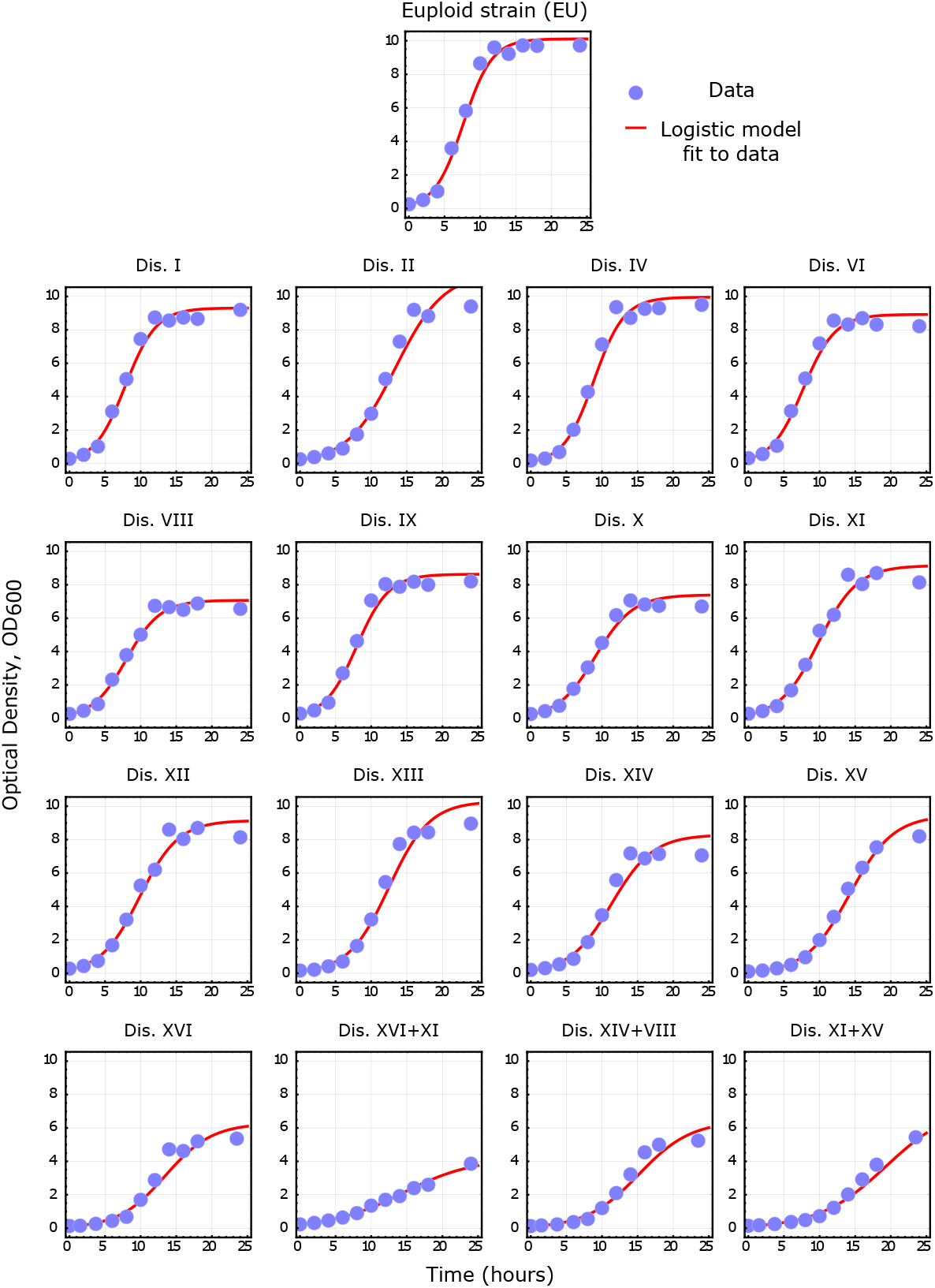
Growth rates of aneuploid strains from ref. [19] are inferred with a logistic fit of growth curves. Shown here are growth curves (Optical Density vs time) of the aneuploid strain and of the euploid strains with the same genetic background taken form Torres at al. The experimental data (circles) are shown together with a logistic fit (solid red lines). Model parameters were inferred with a Bayesian framework (see Material and Methods). Values of the posterior mean values are reported in Table S2.

**Figure S5:**
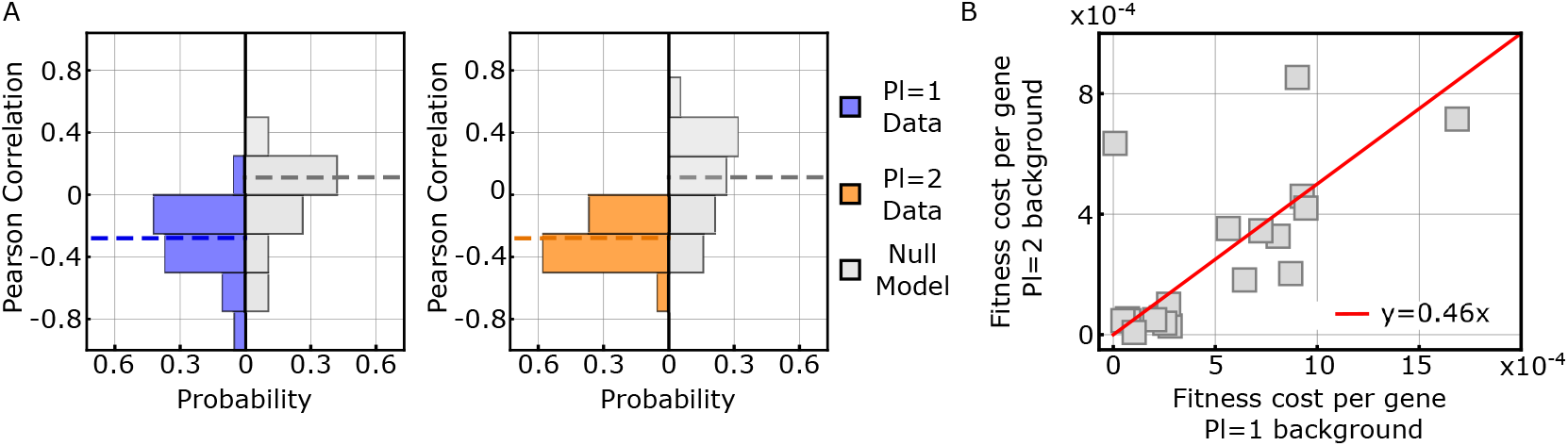
Linear negative correlations between growth rates and number of genes in exceeding chromosomes of aneuploid strains are coherently observed in all growth conditions. A: Histograms of the values of the Pearson’s correlation coefficients of aneuploidy strains with a Ploidy=1 (Left) and a Ploidy=2 background (Right, for the data-set from ref. [9]. In each condition, the Pearson coefficient was evaluated between the values of strain growth rates and the corresponding total number of genes contained in the aneuploidy chromosomes. The null model was obtained with a randomization test, where, in each conditions, growth rates and number of exceeding genes were randomly shuffled and the Pearson correlation coefficient was evaluated for the shuffled data. Dashed lines mark the mean value of each histogram. The difference between the null model distribution and the corresponding distributions of Pl=1 and Pl=2 background are statistically significant (*p*_*val*_ = 0.00003, 0.005 respectively, Mann Whitney U test). B: Scatter plot for the fitness cost per gene (*c*_0_) evaluated for Pl=1 and Pl=2 strains (each data-point correspond to the same condition). The values of the fitness cost display a statistically significant linear correlation (Pearson Correaltion coefficient 0.68, *p*_*val*_ *<* 0.002). The red line show the best linear fit of the data-points.

**Figure S6:**
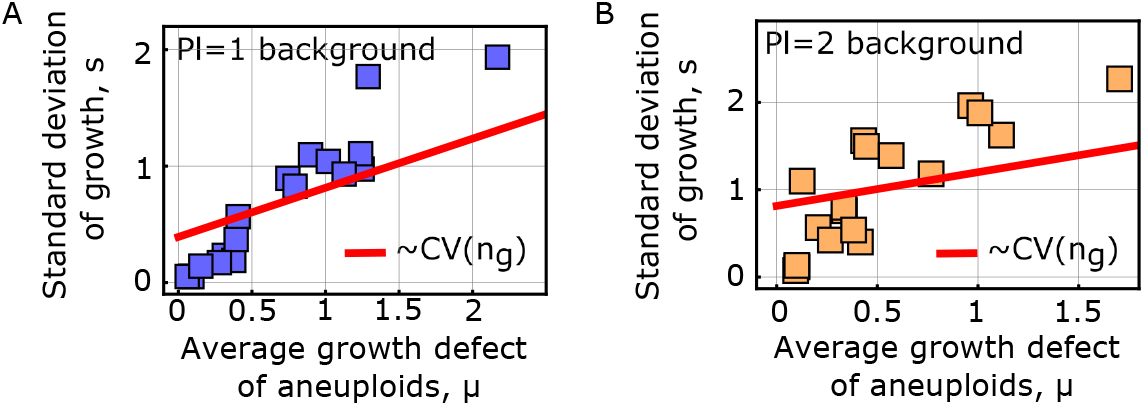
The fitness cost model predicts a linear relationship between the average fitness defect of aneuploids and standard deviation of growth rates, and explains about 80% of the observed dispersion. A and B: Model predictions capture large-scale phenotype data from ref. [28]. The scatter plots of the average growth rate defect (*μ*) of an aneuploid strain across several conditions (environments and stresses) versus its standard deviation across the same set of conditions (*s*). Each square represents a strain of the aneuploid collection of ref. [9] (see Materials and Methods for description of the data-set). Panel A refers to a haploid background, whereas panel B refers to a diploid background. Our form of the fitness cost predicts a linear relation between *μ* and *s*, with a slope equal to the coefficient of variation (CV) of the distribution of the number of excess genes (*n*_*g*_) contained in the aneuploid chromosomes across the set of strains (see Materials and Methods). The red lines show that the this model components alone can describe the observed linear trend and explain the data only partially, with values of the *R*^2^ statistics: 0.84, 0.80 for panel A and B respectively,

**Figure S7:**
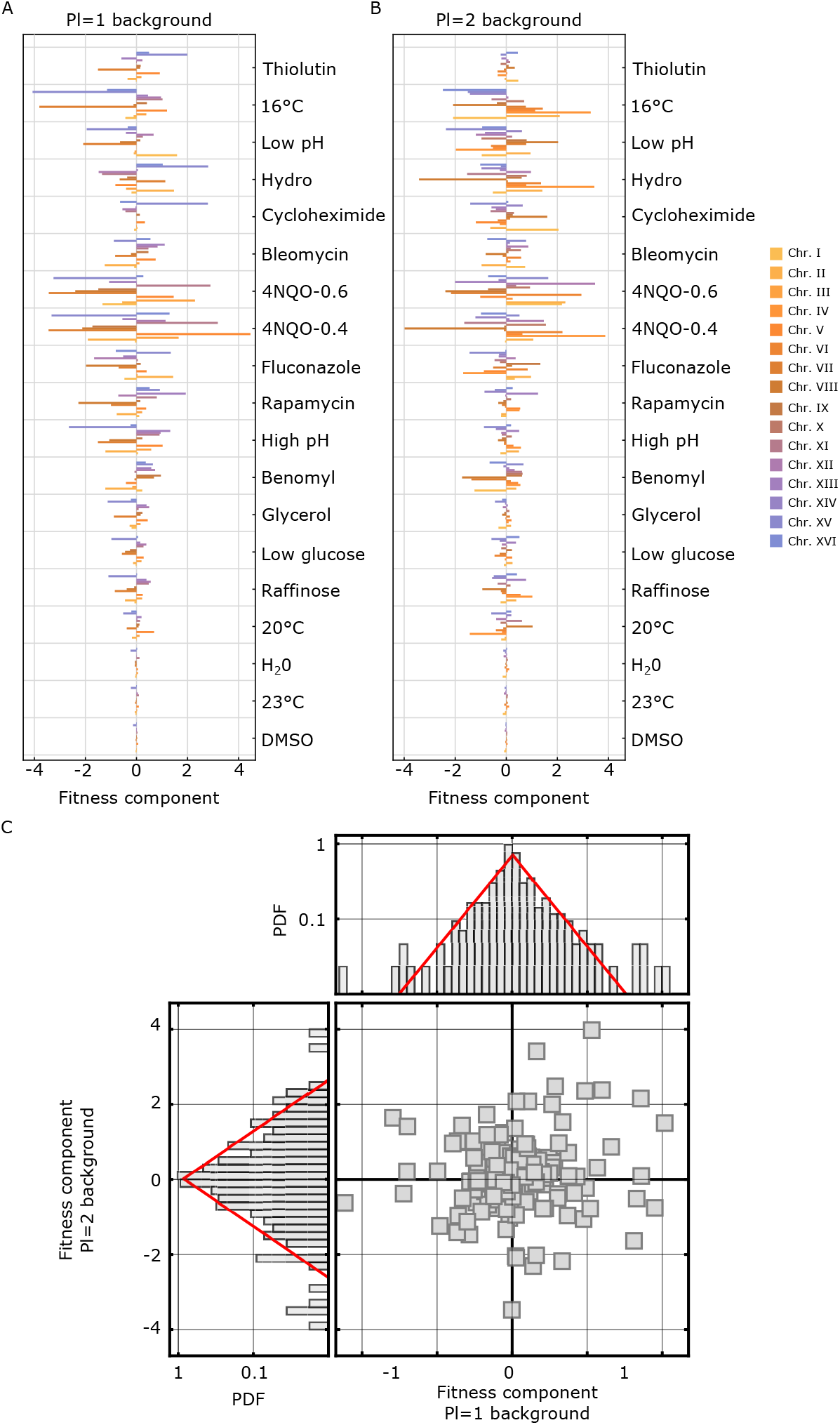
Chromosome fitness components of aneuploid strains can be inferred from measured growth rates. The inferred fitness component of A, a disomic chromosome in a ploidy=1 background, and B, a trisomic chromosome in a ploidy=2 background, is shwon here as a function of the stress condition in which growth rates were evaluated (shown in the x axis). In both the two panels, the chromosome-specific component is shown with a bar plot color coded according to the legend shown on the right. Chromosome fitness components were inferred from growth rates obtained from [9]. Details about the inference are given in Materials and Methods.B. Statistics of the inferred values of the selection coefficients of aneuploidy chromosomes, inferred from [9],and evaluated across several stress conditions. The central panel shows the scatter plot for the values of the chromosomal selection coefficient evaluated in strains with ploidy=1 background vs ploidy=2 background (each square correspond to the same condition and the same chromosome). The data does not show statstically significant correlation (Pearson’s r=0.11). Left and top panels show the probability distribution density of the chromosomal selection coefficients across all conditions and for all chromosomes together for (Top) ploidy=1 and (Left) ploidy=2 background, respectively (gray boxes). The two distributions are in very good agreement with a Laplace distribution (red lines, mean values −0.01, 0.02 and variance 1, 0.7 respectively). Note that in the central panel we report only components that are present in both data-sets.

**Figure S8:**
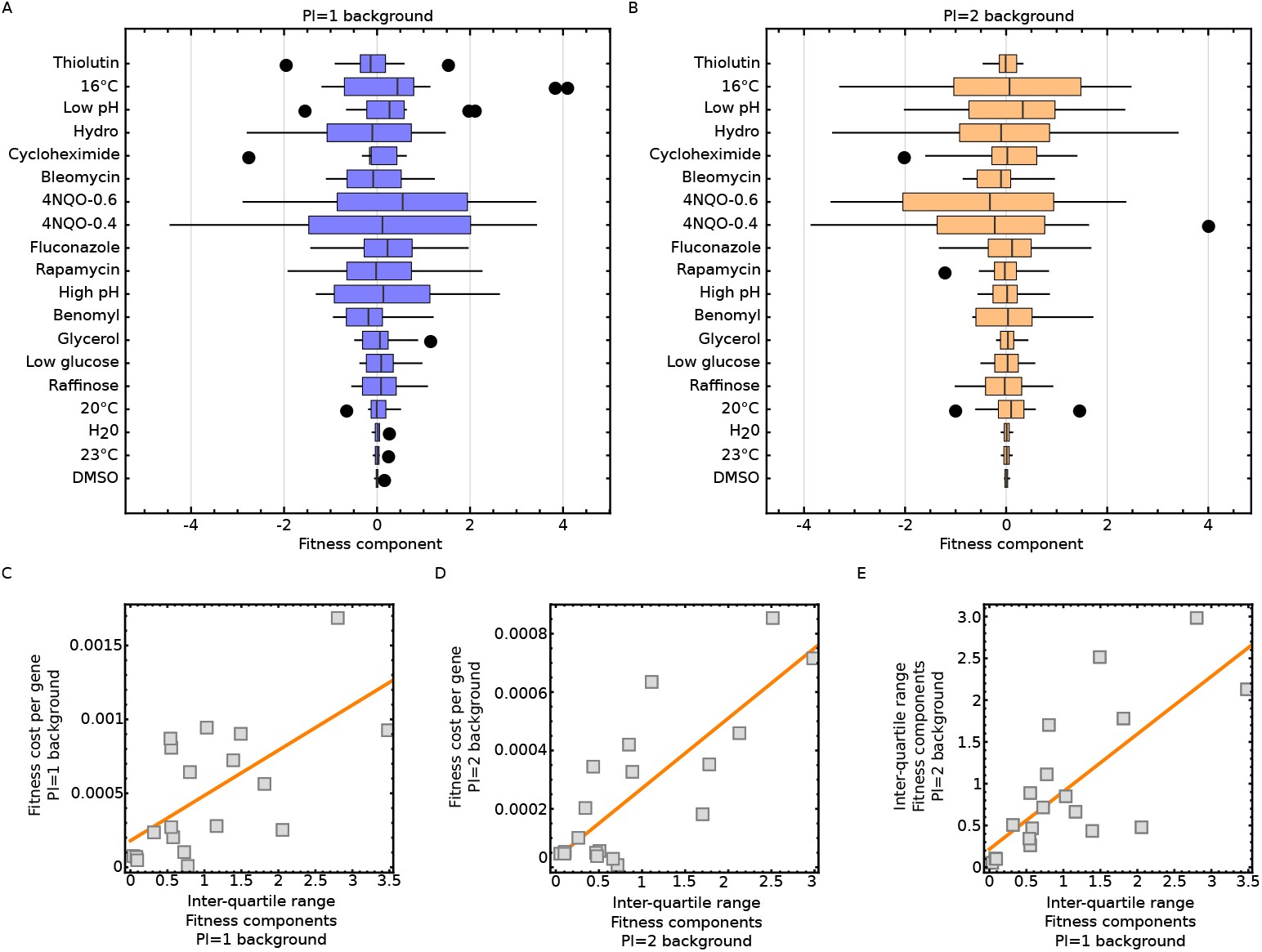
The statistical properties of the distributions of chromosomal fitness components classify environments by harshness. Panels A and B show Box-Whisker plots for the distributions of the inferred values of the chromosome-specific fitness components for each environment. Panel A shows the distributions for ploidy=1 background and panel B shows the distributions for ploidy=2 background. In the boxes, the black line marks the mean value of the distribution, the boxes make the lower (Q1) and upper (Q3) quartile. Whiskers mark the minumum-maximum values and circles mark outliers (points beyond the inter-quartile range from the edge of the box). C: Scatter plot of the inter-quartile ranges of the distributions shown in A *vs* fitness cost per gene in the ploidy=1 background, supporting the idea that the environment-specific inter-chromosome variability of fitness effects is a proxy of environmental harshness. D: Scatter plot for the inter-quartile ranges of the distributions shown in B *vs* fitness cost per gene in the pl=2 background. E: Scatter plot between the inter-quartile ranges of the distributions for ploidy=1 and polidy=2 backgrounds, showing that the width of the distributions are correlated. In panels C-D-E we observe a statistically significant linear correlation coefficient, with Pearson’s r values= (0.8, 0.7, 0.6) and p-val= (0.00004, 0.0003, 0.003) respectively.

**Figure S9:**
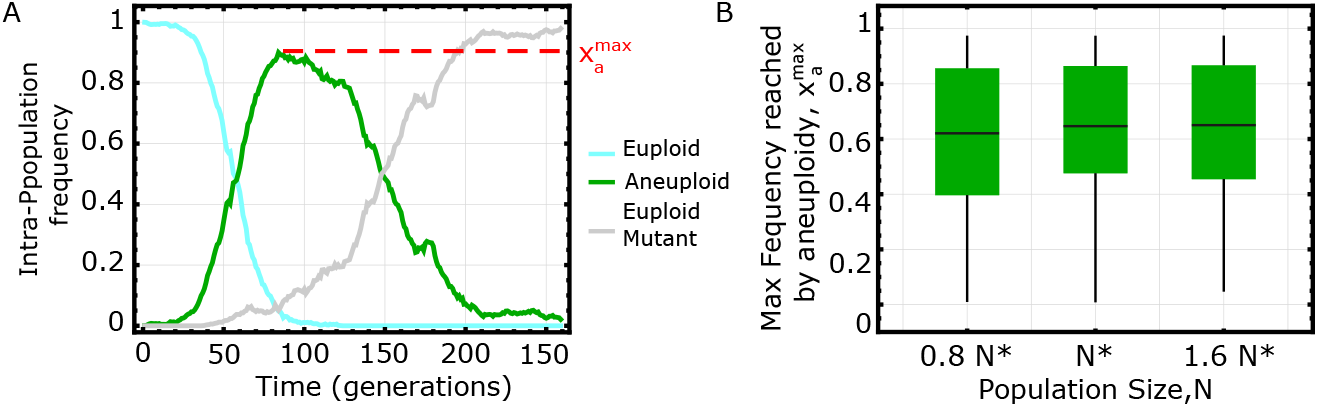
Clonal Interference (C.I.) effects can be mistaken for a loss of aneuploidy resulting from karyotype instability. We show here the results of numerical simulations of our evolutionary model (cfr. Material and Methods for details about the simulations)to illustrate the model predictions in the CI regime. In (A) we show one simulation instance, to illustrate the typical dynamics of the intra-population frequencies of euploid individuals (cyan), aneuploid individuals (green) and of euploid mutants (gray). The dynamics displays two distinct phases: (i) a rise in frequency of aneuploid individuals up to a maximum value 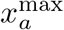, followed by (ii) a decline and subsequent elimination of aneuploidy from the population, because of the emergence of an euploid mutant. B. Probability distribution of the maximal frequency reached by aneyploidy before being eliminated from the population because of CI *vs* the value of the population size used in the simulation. Probabillity distributions are shown with box-whisker plots (black line: mean value, box: inter-quartile range, fences: max and min values). Here, the population size is expressed in terms of the critical population size *N* ^*^ (Eq.31 in S.I.); evolutionary experiments performed with an effective population size *N N* ^*^ are governed by effects C.I. and have a high probability (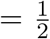 for *N* = *N* ^*^) to display two distinct phases. Taken together, these results show that, when CI is very likely to take place (i.e., for *N N* ^*^, cfr S.I.) since the anueploids can reach a substantial maximum frequency 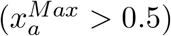, this dynamics could be wrongly interpreted as a fixation of aneuploids followed by loss of the extra chromosome. Model parameters used in the simulations: (i) selection coefficients *σ*_*b*_ = 0.17 gen^−1^, *σ*_*c*_ = 0.05 gen^−1^),(ii) mutation rates *μ*_*a*_ = 4 * 10^−4^gen^−1^ and*μ*_*m*_ = 2 * 10^−5^gen^−1^, (iii) population size *N* = 2000 (Panel A), 1500,2000,2500 (Panel B). With this set of model parameters, the critical population size (Eq.31 in S.I.) takes the value *N* ^*^ = 1983 ≃ 2000.

**Figure S10:**
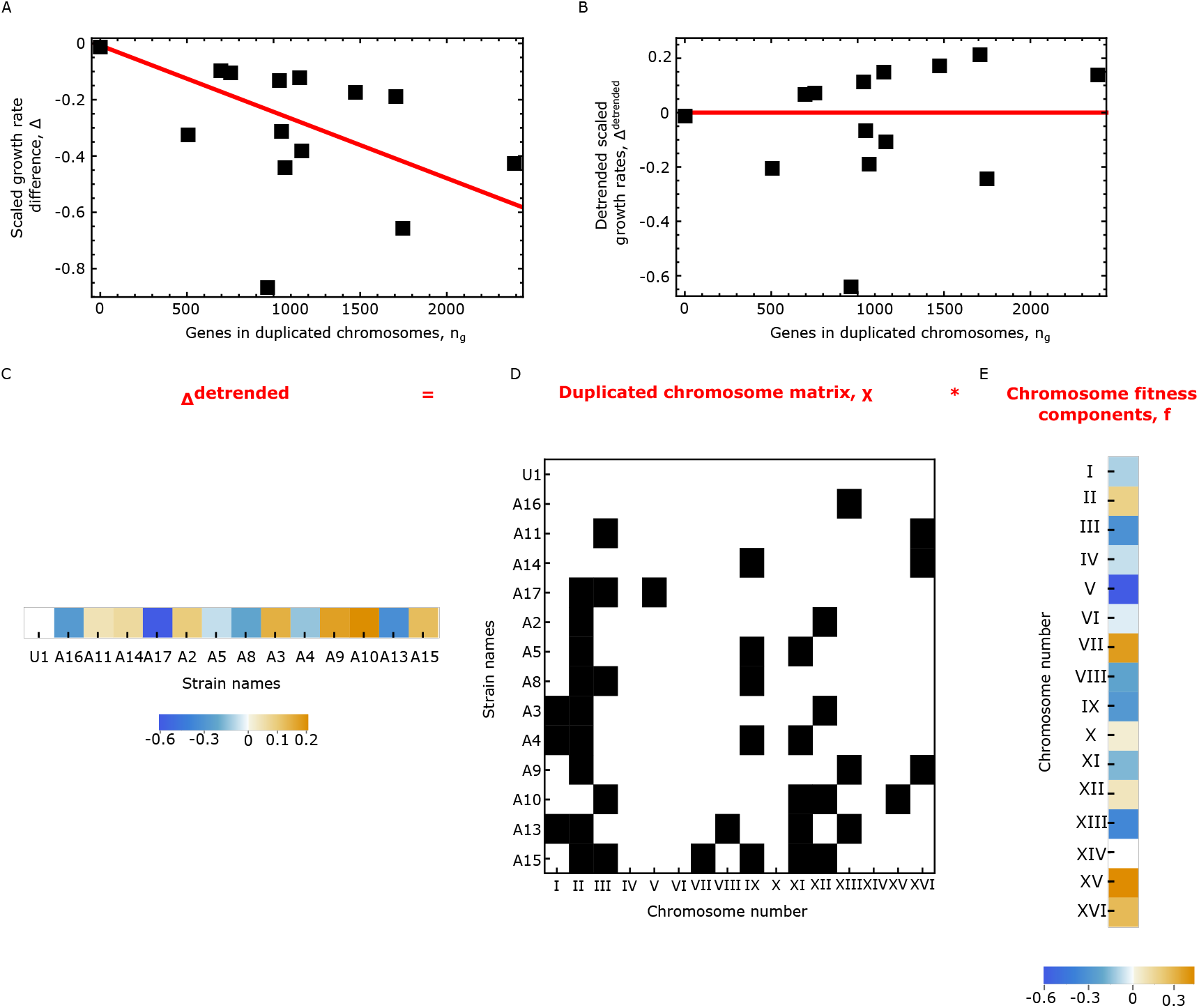
Detailed procedure of the inference of chromosomal fitness components from scaled growth rate. We show here a detailed example of the algorithmic and mathematical steps involved in the inference of the fitness components from scaled growth rates from ref. [9]. In this example, we show growth data evaluated at 20°C, for strains with for pl=1 background. Panel A shows the values of the scaled growth rate differences Δ_*s*_ (computed according to Eq. 19) *vs* the number of duplicated genes contained in the corresponding strain 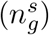. The red line shows the linear decreasing trend for this data, which, in our model, is related to the average fitness cost of a gene contained in a duplicated chromosome. The first step of our inference removes this (genome-wide) linear trend, by detrending the data (according to Eq.23). The result of this detrending procedure is shown in panel B, the red line now showing the absence of a residual trend. The set of detrendend data of all the considered strains can be mathematically represented as a vector 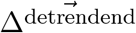, as shown in panel C, where numerical values are now color coded (legend at the bottom of the panel). To infer the chromosome-specific fitness components we decompose this set of values 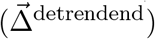 into the product of the duplicated chromosomes *χ* and the vector of the chromosomal fitness components 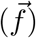. The matrix *χ*, shown in panel D, specifies the set of duplicated chromosomes in a given strain; here duplicated chromosomes are shown in black (corresponding to a numerical value =1) while non-duplicated chromosomes are shown in white (corresponding to a numerical value =0). The vector of the fitness components specifies the contribution of each chromosome to the observed values of the detrended growth rates 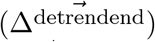 and is computed according to Eq. 23. Note that the algebraic relation 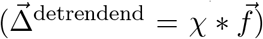 implies that the detrended growth rate of a given strain is the sum of the fitness components of its duplicated chromosomes only.

**Table S1:**
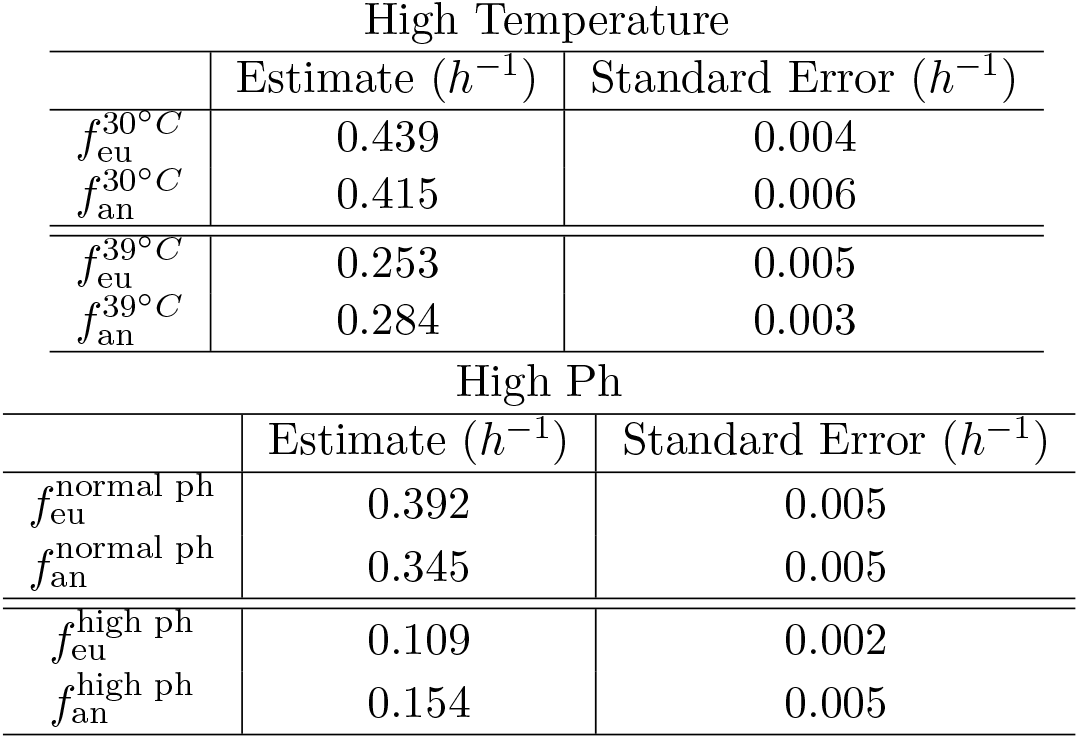
Inferred Values of the growth rates of the aneuploid and euploid strain from ref. [20].

| High Temperature |  |  |
| --- | --- | --- |
| | Estimate ($h^{-1}$) | Standard Error ($h^{-1}$) |
| $f_{eu}^{30^{\circ}C}$ | 0.439 | 0.004 |
| $f_{an}^{30^{\circ}C}$ | 0.415 | 0.006 |
| $f_{eu}^{39^{\circ}C}$ | 0.253 | 0.005 |
| $f_{an}^{39^{\circ}C}$ | 0.284 | 0.003 |
| High Ph |  |  |
| | Estimate ($h^{-1}$) | Standard Error ($h^{-1}$) |
| $f_{eu}^{normal\ ph}$ | 0.392 | 0.005 |
| $f_{an}^{normal\ ph}$ | 0.345 | 0.005 |
| $f_{eu}^{high\ ph}$ | 0.109 | 0.002 |
| $f_{an}^{high\ ph}$ | 0.154 | 0.005 |

**Table S2:** Posterior mean values of the model parameters for the logistic fit 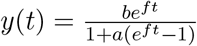 of the growth data from ref. [19].

| Strain | # exceeding genes | $f$ ($h^{-1}$) | a | b | inverse variance $1/\sigma_Y^2$ |
| --- | --- | --- | --- | --- | --- |
| Eu | 0 | 0.5 | 0.022 | 0.22 | 38 |
| Dis. I | 117 | 0.46 | 0.026 | 0.25 | 48 |
| Dis. II | 456 | 0.46 | 0.025 | 0.24 | 48 |
| Dis. IV | 836 | 0.2 | 0.053 | 0.22 | 169 |
| Dis. VI | 139 | 0.29 | 0.019 | 0.22 | 35 |
| Dis. VIII | 321 | 0.45 | 0.031 | 0.28 | 43 |
| Dis. IX | 241 | 0.47 | 0.015 | 0.15 | 35 |
| Dis. X | 398 | 0.44 | 0.028 | 0.24 | 40 |
| Dis. XI | 348 | 0.42 | 0.033 | 0.24 | 51 |
| Dis. XII | 578 | 0.34 | 0.02 | 0.17 | 29 |
| Dis. XIII | 505 | 0.37 | 0.026 | 0.24 | 48 |
| Dis. XIV | 435 | 0.38 | 0.031 | 0.23 | 61 |
| Dis. XV | 597 | 0.33 | 0.0095 | 0.09 | 55 |
| Dis. XVI | 511 | 0.35 | 0.012 | 0.13 | 25. |
| Dis. XVI+XI | 859 | 0.27 | 0.016 | 0.1 | 19 |
| Dis. XIV+VIII | 756 | 0.3 | 0.016 | 0.1 | 17 |
| Dis. XI+XV | 945 | 0.21 | 0.016 | 0.12 | 33 |

**Table S3:**
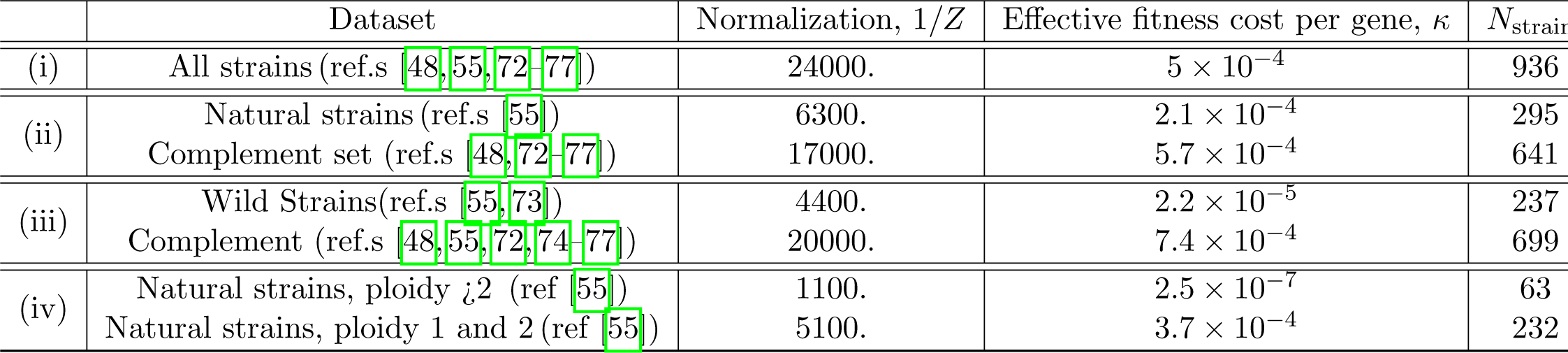
Best-fit model parameters for the aneuploid strains abundances data ([21]).*Numerical values of the model parameters for the fit of Eq.**??** to aneuploidy strains abundances data. In (i) we considered the full data-set presented in ref. ([21], where authors aggregated data for aneuploidy strains from 8 independent studies. In (ii) we split the data-set into two subsets, the subset of “natural strains” from ref. [55] and its complement set, in order to evaluate differences between model parameters for the two subsets. Similarly, in (iii) we compared the set of “wild strains” from ref. [21] and its complement set. In (iv) we focused on the subset of natural strains only, which we further partitioned into aneuploid strains with a ploidy¿2 background and strains with ploidy=1 and ploidy=2 background.*

## Supplementary Information Appendix

### Detailed derivation of the analytical results of the evolutionary model

#### Onset of Aneuploidy

We focus on the waiting times for the emergence of a successful mutant (defined as the mutant that will eventually reach fixation). The two times, denoted as *t*_*a*_ and *t*_*m*_ for the aneuploid and the euploid mutant respectively, are stochastic variables, with expected values equal to the inverse of the fixation rates (*τ*_*a,m*_≡ *(t*_*a,m*_) = 1*/λ*_*a,m*_), and with exponential probability distribution

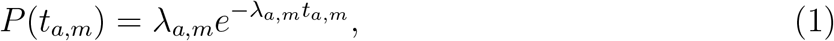

The statistics of the fastest emerging mutant can described by the difference of the two times

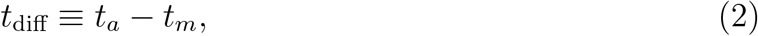

whose probability density reads

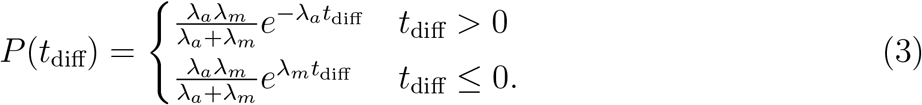

The problem of computing the probability for aneuploidy to reach fixation is equivalent to computing the probability for the time difference to be negative (*t*_diff_ *<* 0). Clonal Interference effects are captured by the extended condition 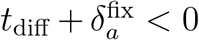, where 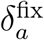 is the effective time to fixation of an aneuploid mutant (see next paragraph). This leads to the expression

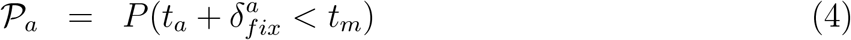

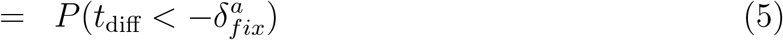

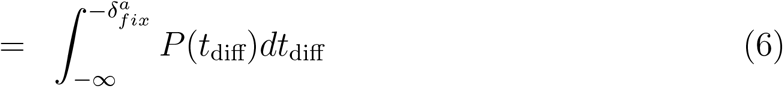

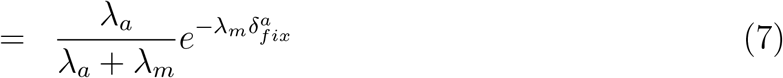

The waiting time until the emergence of the fastest successful mutant can also be investigated through the statistical properties of the minimum of the two waiting times, *t*_min_ ≡ min(*t*_*a*_, *t*_*b*_)

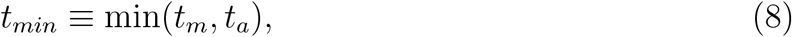

whose expected value reads

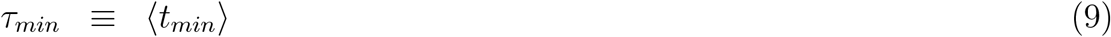

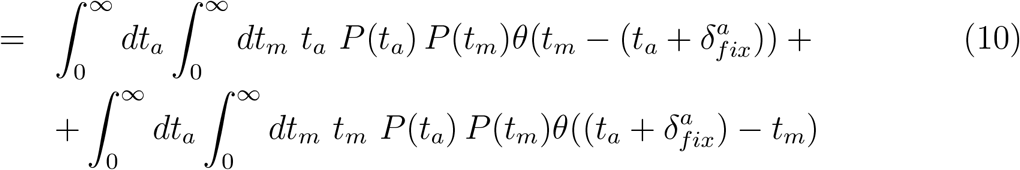

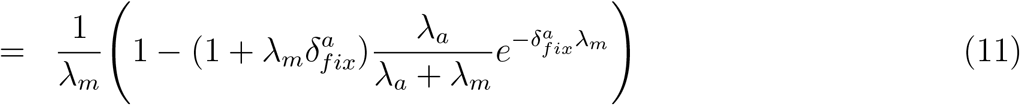

#### Effective time to fixation of the aneuploid mutant

In order to compute the effective time where Clonal Interference effects can take place, we follow the same argument presented in [2, 3]. We denote with *x*(*t*) the intra-population frequency of the wild-type (not-mutated and euploid) strain, and with *σ*_*a*_ *>* 0 the selection coefficient of the aneuploid mutatant, evaluated w.r.t the wild type. The effective population size is *N*. In the deterministic limit, i.e., neglecting genetic drift effects, the decline in frequency of the wild-type strain is described by the logistic equation

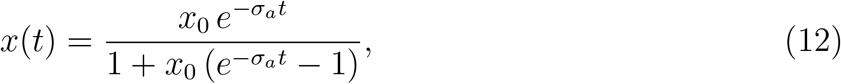

where *x*_0_ ≡ *x*(*t* = 0). Interference effects can take place during the deterministic dynamics, while genetic drift effects dominate the dynamics for *x*(*t*) ≥ *x*_0_ ≃ 1 and *x*(*t*) ≤1 −*x*_*f*_.

Following the argument presented in [3] we use the boundaries where (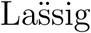 boundaries)

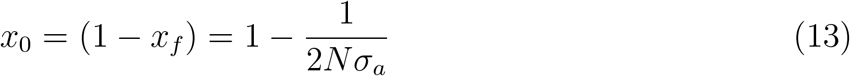

Hence, the time interval during which an interfering mutation can arise is given by

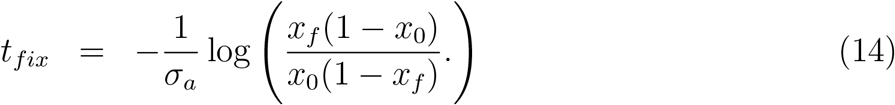

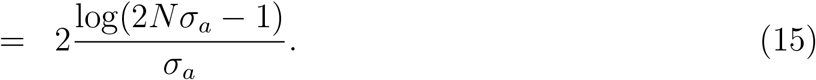

Euploid mutants that can emerge from wild type strains with a mutation rate *μ*_*m*_ during this time interval have expected number

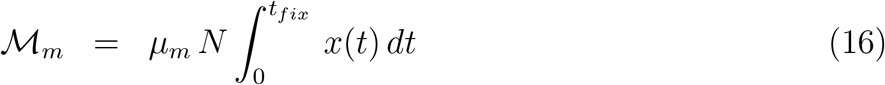

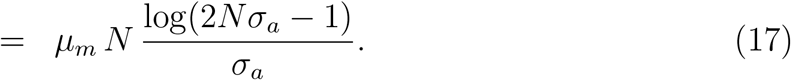

Finally, the number of mutations that can interfere with the aneuploid mutant are given by

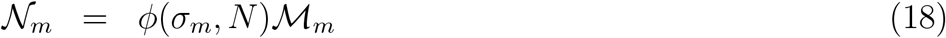

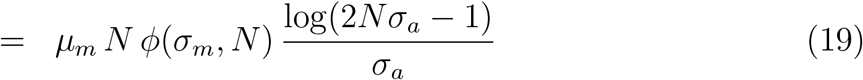

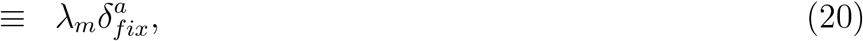

where *λ*_*m*_ = *μ*_*m*_*Nϕ*(*σ*_*m*_, *N*) is the fixation rate of the euploid mutant and we have defined the effective time to fixation as

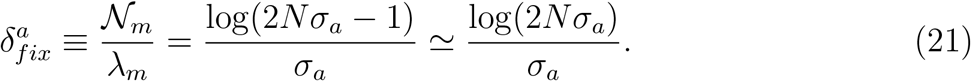

#### Critical value of the beneficial selection coefficient 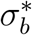

We find the critical value of the beneficial selection coefficient by solving the equation

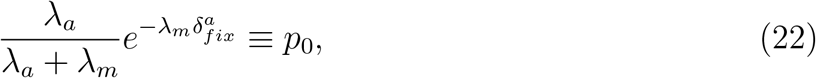

where 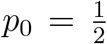. We solve this equation using the Haldane’s Formula [4] for the fixation probability

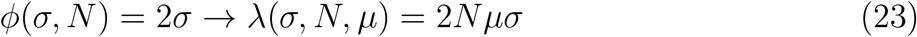

and we consider the following dynamical regimes

i. *No Clonal Interference*. This regime corresponds to the mathematical condition 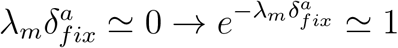, i.e., the expected number of euploid mutants interfering with the fixation dynamics of the aneuploid mutant are negligible. In this regime, Eq.22 reads

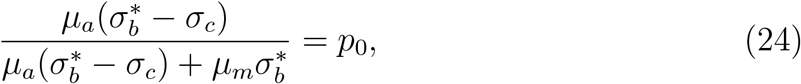

and the solution reads

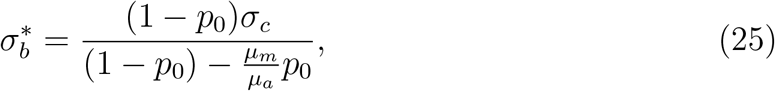

where 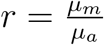.
ii. *In the Clonal Interference regime*. In this regime we find an approximated solution to Eq.22 by considering the first order expansion 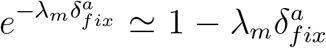. With this approximation, Eq.22 reads

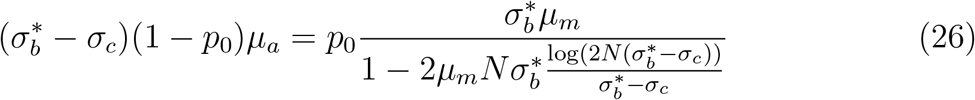

by approximating 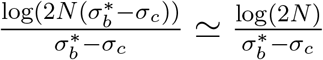 we get the expression

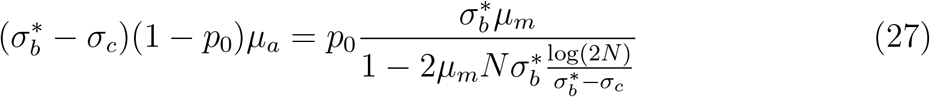

with the explicit analytic solution

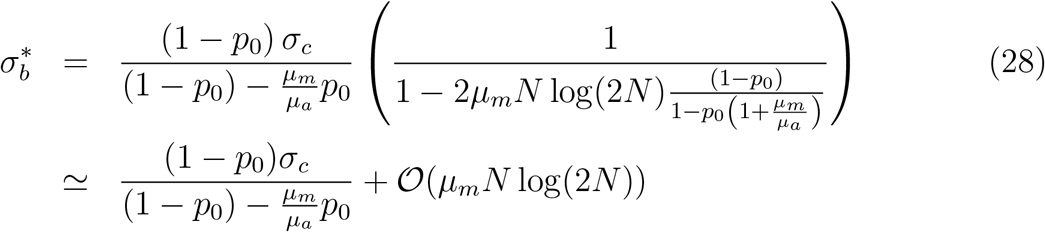

Hence, the general solution that recapitulates both dynamical regime is obtained for *p*_0_ = 1*/*2 and reads

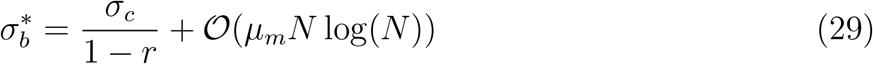

where *r* = *μ*_*m*_*/μ*_*a*_.

### High Ph experiment data from ref. [5]

In ref. [5], Yona and coworkers investigated an evolved strain obtained from an experiment presented in a previous publication [6], in which a haploid S. cerevisiae was evolved under high PH (8.6), using a transfer protocol similar to the one used for the High T experiment. In such experiment (one replicate only) the duplication of chromosome V (trisomy) was found to have reached fixation after about ∼150 generations; the duplication of this chromosome shown to confer a beneficial effect in response to the applied stress.

Growth curves relative to this strains (aneuploid and euploid) were obtained from spores produced at the end of the experiments, and were used here to infer growth rates. It should be noted that these two strains, namely the euploid reference and aneuploid strains, contained 4 point mutations that emerged during the experiment and were not present in the euploid strain used at the beginning of the experiment. This set of mutations could possibly confer additional advantage to the high PH [6].

In this context we neglected the effect of these mutations, and we showed that the evolutionary dynamics leading to the emergence of resistance to High-PH can be explained by the emerging dynamics of aneuploidy alone. It should be noted, in particular, that the difference in fitness between the initial euploid strain and the aneuploid strain is higher than the one observed (euploid vs aneuploid, both of them with the set of adaptive mutations). Our estimate of *σ*_*b*_− *σ*_*c*_ is therefore a conservative one, since it quantifies the minimal fitness advantage conferred by the chromosome gain alone.

### Relaxing the assumption for the selection coefficient of euploid mutant

This section reviews the model assumption used throughout the paper that concerns the selection coefficient of the euploid mutant. We show how our model maps to more complex scenarios within the same model formalism. More specifically, we have assumed that, in response to external stress, an euploid individual developing point mutations can gain a fitness benefit equal to the maximal fitness gain attained by doubling the expression of the target gene

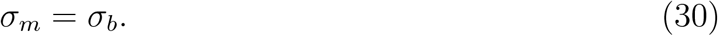

Using this assumption, we have considered a conservative scenario, i.e., the one where the emergence of aneuploidy is mostly suppressed. However, in a more general scenario, one could focus on point mutations that alter gene expression to a lower degree, hence, investigate the competition between aneuploid individuals and euploid mutants with a lower selection coefficient

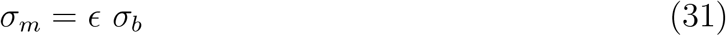

with 0 < *ϵ* ≤ 1 and with a fixation rate

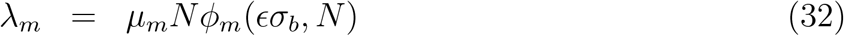

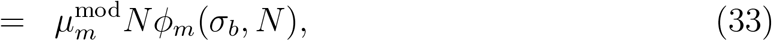

where *ϕ*_*m*_(*σ*_*m*_, *N*) is the fixation probability of the euploid mutant in a population with *N* individuals and we have definied the modified mutation rate

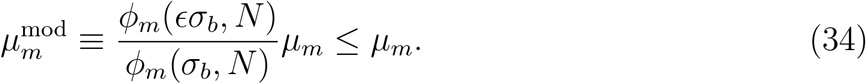

In other words, from Eq.s (33,34), one can deduce that a model with a selection coefficient *σ*_*m*_ = *fσ*_*b*_ is equivalent to a model with *σ*_*m*_ = *σ*_*b*_ and a reduced mutation rate 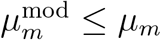. Hence, relaxing the assumption Eq.30 corresponds to considering an effective mutation rate 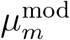 which could be lower than the per-base spontaneous mutation rate. Of note, when considering the Haldane’s Formula [4] for the fixation probability, one has 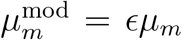. Considering the comparison of our model with data from the evolutionary experiment [5], we find that the model prediction is in agreement with the experimental data also for values of mutational rates lower than the spontaneous per-nucleotide error rate *μ*_*m*_ ≤ *μ*_spont._ = 1.7 * 10^−10^gen^−1^ (cfr. Fig.2, S2, S3). Hence, our model would also support a scenario where the euploid mutant is characterized by a selection coefficient *σ*_*m*_ ≤ *σ*_*b*_.

### Loss of aneuploidy in the clonal interference regime

This section addresses the quantitative predictions of our model that are key for the interpretation of evolutionary experiments aimed at investigating the fate of aneuploid individuals, i.e., testing whether or not this karyotype state is genetically stable. In experimental setups akin to that used in ref. [5], the loss of aneuploidy in the long term could be explained by two alternative scenarios: (i) loss of the aneuploidy and subsequent gain of the point mutations or (ii) elimination of aneuploid individuals resulting from clonal interference effects (the euploid point mutation emerges before the complete fixation of aneuploidy, and outcompetes it). Scenario (i) would imply that aneuploid individuals are not stable (the extra chromosome is lost) while scenario (ii) does not exclude this possibility (in this case aneuploid individuals did not lose the extra chromosome). In the following, we will give quantitative conditions for scenario (ii) to be observed in experimental setups akin to ref. [5].

To illustrate this point, we consider a conservative scenario where the emergence of aneuploidy would be very likely if no Clonal Inferference (C.I.) effects were present, and corresponds to the mathematical condition

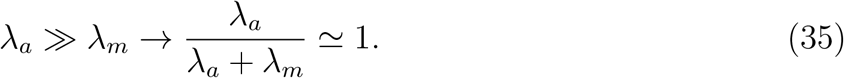

This model condition is fulfilled by the model parameters estimated for the experiment of ref. [5] (for which *σ*_*b*_ = 0.17gen^−1^ and *σ*_*b*_ = 0.12gen^−1^ and using realistic values of the mutational rates *μ*_*a*_≃ 10^−6^gen^−1^ and *μ*_*m*_≤ 10^−8^gen^−1^). Hence, this regime describes a realistic scenario for evolutionary experiments akin to [5].

In this case, the probability to develop aneuploidy reads

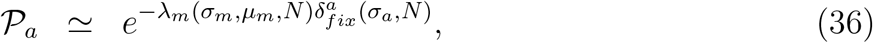

and predicts C.I. effects to dominate the dynamics and substantially reduce the probability to develop aneuploidy for experiments carried with an effective population sizes larger than a critical value

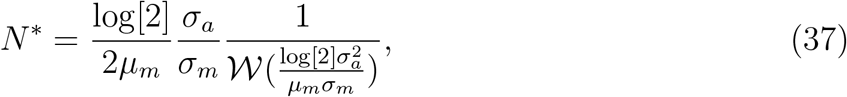

where 𝒲 (*x*) is the Lambert-W function (note that Eq.(37) is the solution to the equation 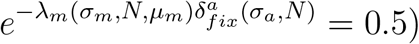.

The dynamics predicted by our model in this regime for a typical evolutionary experiment (akin to ref. [5]) is shown in Fig. S9 A. When the effective population size used during the experiment is close to the critical value (*N* ≃*N* ^*^) we find that aneuploids can reach a high intra-population frequency before being replaced by the euploid point mutation (Fig. S9 B.). Hence, in this regime, the prediction of our model is that the loss of aneuploid individuals from the population could be mistaken as a sign of genomic instability while, instead, it is an indirect effect caused by C.I.

## Notes

### Competing Interest Statement

The authors have declared no competing interest.

### Summary of Updates

Introduction and Conclusion updated to (i) clarify the interpretation of the model concerning the fitness cost per gene and (ii) to illustrate how our model can be used to investigate Clonal Interference effect in evolutionary experiments involving the emergence of anueploidy strains; Supplemental files uploaded

## References

1. Douglas Hanahan and Robert A Weinberg. Hallmarks of cancer: the next generation. cell, 144(5):646–674, 2011.

2. Laurent Sansregret and Charles Swanton. The role of aneuploidy in cancer evolution. Cold Spring Harbor perspectives in medicine, 7(1):a028373, 2017.

3. Alison M Taylor, Juliann Shih, Gavin Ha, Galen F Gao, Xiaoyang Zhang, Ashton C Berger, Steven E Schumacher, Chen Wang, Hai Hu, Jianfang Liu, et al. Genomic and functional approaches to understanding cancer aneuploidy. Cancer cell, 33(4):676–689, 2018.

4. Richard D Kolodner, Don W Cleveland, and Christopher D Putnam. Aneuploidy drives a mutator phenotype in cancer. Science, 333(6045):942–943, 2011.

5. Rustelle Janse van Vuuren, Michelle H Visagie, Anne E Theron, and Annie M Joubert. Antimitotic drugs in the treatment of cancer. Cancer chemotherapy and pharmacology, 76(6):1101– 1112, 2015.

6. Mattia Pavani, Paolo Bonaiuti, Elena Chiroli, Fridolin Gross, Federica Natali, Francesca Macaluso, Ádám Póti, Sebastiano Pasqualato, Zoltán Farkas, Simone Pompei, et al. Epistasis, aneuploidy, and functional mutations underlie evolution of resistance to induced microtubule depolymerization. The EMBO Journal, page e108225, 2021.

7. Hung-Ji Tsai, Anjali R Nelliat, Mohammad Ikbal Choudhury, Andrei Kucharavy, William D Bradford, Malcolm E Cook, Jisoo Kim, Devin B Mair, Sean X Sun, Michael C Schatz, et al. Hypo-osmotic-like stress underlies general cellular defects of aneuploidy. Nature, 570(7759):117– 121, 2019.

8. Allegra Terhorst, Arzu Sandikci, Abigail Keller, Charles A Whittaker, Maitreya J Dunham, and Angelika Amon. The environmental stress response causes ribosome loss in aneuploid yeast cells. Proceedings of the National Academy of Sciences, 117(29):17031–17040, 2020.

9. Norman Pavelka, Giulia Rancati, Jin Zhu, William D Bradford, Anita Saraf, Laurence Florens, Brian W Sanderson, Gaye L Hattem, and Rong Li. Aneuploidy confers quantitative proteome changes and phenotypic variation in budding yeast. Nature, 468(7321):321–325, 2010.

10. Jason M Sheltzer and Angelika Amon. The aneuploidy paradox: costs and benefits of an incorrect karyotype. Trends in Genetics, 27(11):446–453, 2011.

11. Anna B Sunshine, Celia Payen, Giang T Ong, Ivan Liachko, Kean Ming Tan, and Maitreya J Dunham. The fitness consequences of aneuploidy are driven by condition-dependent gene effects. PLoS Biol, 13(5):e1002155, 2015.

12. Dawn A Thompson, Michael M Desai, and Andrew W Murray. Ploidy controls the success of mutators and nature of mutations during budding yeast evolution. Current Biology, 16(16):1581– 1590, 2006.

13. Aleeza C Gerstein. Mutational effects depend on ploidy level: all else is not equal. Biology letters, 9(1):20120614, 2013.

14. Kristoffer Krogerus, Mikko Arvas, Matteo De Chiara, Frederico Magalhães, Laura Mattinen, Merja Oja, Virve Vidgren, Jia-Xing Yue, Gianni Liti, and Brian Gibson. Ploidy influences the functional attributes of de novo lager yeast hybrids. Applied microbiology and biotechnology, 100(16):7203–7222, 2016.

15. Clifford Zeyl. Experimental studies of ploidy evolution in yeast. FEMS microbiology letters, 233(2):187–192, 2004.

16. Aleeza C Gerstein and Sarah P Otto. Ploidy and the causes of genomic evolution. Journal of Heredity, 100(5):571–581, 2009.

17. Jenny M McLaughlan, Gianni Liti, Sarah Sharp, Agnieszka Maslowska, and Edward J Louis. Apparent ploidy effects on silencing are post-transcriptional at hml and telomeres in saccharomyces cerevisiae. PLoS One, 7(7):e39044, 2012.

18. Christopher W Bakerlee, Angela M Phillips, Alex N Nguyen Ba, and Michael M Desai. Dynamics and variability in the pleiotropic effects of adaptation in laboratory budding yeast populations. Elife, 10:e70918, 2021.

19. Eduardo M Torres, Tanya Sokolsky, Cheryl M Tucker, Leon Y Chan, Monica Boselli, Maitreya J Dunham, and Angelika Amon. Effects of aneuploidy on cellular physiology and cell division in haploid yeast. science, 317(5840):916–924, 2007.

20. Avihu H Yona, Yair S Manor, Rebecca H Herbst, Gal H Romano, Amir Mitchell, Martin Kupiec, Yitzhak Pilpel, and Orna Dahan. Chromosomal duplication is a transient evolutionary solution to stress. Proceedings of the National Academy of Sciences, 109(51):21010–21015, 2012.

21. Ciaran Gilchrist and Rike Stelkens. Aneuploidy in yeast: Segregation error or adaptation mechanism? Yeast, 36(9):525–539, 2019.

22. Jin Zhu, Hung-Ji Tsai, Molly R Gordon, and Rong Li. Cellular stress associated with aneuploidy. Developmental cell, 44(4):420–431, 2018.

23. Sunyoung Hwang, Paola Cavaliere, Rui Li, Lihua Julie Zhu, Noah Dephoure, and Eduardo M Torres. Consequences of aneuploidy in human fibroblasts with trisomy 21. Proceedings of the National Academy of Sciences, 118(6), 2021.

24. Bret R Williams, Vineet R Prabhu, Karen E Hunter, Christina M Glazier, Charles A Whittaker, David E Housman, and Angelika Amon. Aneuploidy affects proliferation and spontaneous immortalization in mammalian cells. Science, 322(5902):703–709, 2008.

25. Silvia Stingele, Gabriele Stoehr, Karolina Peplowska, Jürgen Cox, Matthias Mann, and Zuzana Storchova. Global analysis of genome, transcriptome and proteome reveals the response to aneuploidy in human cells. Molecular systems biology, 8(1):608, 2012.

26. Teresa Davoli, Andrew Wei Xu, Kristen E Mengwasser, Laura M Sack, John C Yoon, Peter J Park, and Stephen J Elledge. Cumulative haploinsufficiency and triplosensitivity drive aneuploidy patterns and shape the cancer genome. Cell, 155(4):948–962, 2013.

27. Nicholas A Graham, Aspram Minasyan, Anastasia Lomova, Ashley Cass, Nikolas G Balanis, Michael Friedman, Shawna Chan, Sophie Zhao, Adrian Delgado, James Go, et al. Recurrent patterns of dna copy number alterations in tumors reflect metabolic selection pressures. Molecular systems biology, 13(2):914, 2017.

28. Guangbo Chen, Wahid A Mulla, Andrei Kucharavy, Hung-Ji Tsai, Boris Rubinstein, Juliana Conkright, Scott McCroskey, William D Bradford, Lauren Weems, Jeff S Haug, et al. Targeting the adaptability of heterogeneous aneuploids. Cell, 160(4):771–784, 2015.

29. Yao Li, Arthur Berg, Louie R Wu, Zhong Wang, Gang Chen, and Rongling Wu. Modeling the aneuploidy control of cancer. BMC cancer, 10(1):1–9, 2010.

30. Andrei Kucharavy, Boris Rubinstein, Jin Zhu, and Rong Li. Robustness and evolvability of heterogeneous cell populations. Molecular biology of the cell, 29(11):1400–1409, 2018.

31. Rong Li and Jin Zhu. Effects of aneuploidy on cell behaviour and function. Nature Reviews Molecular Cell Biology, pages 1–16, 2022.

32. Ronald Aylmer Fisher. The genetical theory of natural selection. The Clarendon Press, 1958.

33. Lev Y Yampolsky and Arlin Stoltzfus. Bias in the introduction of variation as an orienting factor in evolution. Evolution & development, 3(2):73–83, 2001.

34. Erik I Svensson and David Berger. The role of mutation bias in adaptive evolution. Trends in ecology & evolution, 34(5):422–434, 2019.

35. Martijn F Schenk, Mark P Zwart, Sungmin Hwang, Philip Ruelens, Edouard Severing, Joachim Krug, and JAGM De Visser. Population size mediates the contribution of high-rate and large-benefit mutations to parallel evolution. Nature Ecology & Evolution, 6(4):439–447, 2022.

36. Motoo Kimura. On the probability of fixation of mutant genes in a population. Genetics, 47(6):713, 1962.

37. Philip J Gerrish and Richard E Lenski. The fate of competing beneficial mutations in an asexual population. Genetica, 102:127–144, 1998.

38. Michael M Desai, Daniel S Fisher, and Andrew W Murray. The speed of evolution and maintenance of variation in asexual populations. Current biology, 17(5):385–394, 2007.

39. Su-Chan Park and Joachim Krug. Clonal interference in large populations. Proceedings of the National Academy of Sciences, 104(46):18135–18140, 2007.

40. Stephan Schiffels, Gergely J SzöllHosi, Ville Mustonen, and Michael Lässig. Emergent neutrality in adaptive asexual evolution. Genetics, 189(4):1361–1375, 2011.

41. Kavita Jain, Joachim Krug, and Su-Chan Park. Evolutionary advantage of small populations on complex fitness landscapes. Evolution: International Journal of Organic Evolution, 65(7):1945– 1955, 2011.

42. Yuan O Zhu, Mark L Siegal, David W Hall, and Dmitri A Petrov. Precise estimates of mutation rate and spectrum in yeast. Proceedings of the National Academy of Sciences, 111(22):E2310– E2318, 2014.

43. Gal Hagit Romano, Yonat Gurvich, Ofer Lavi, Igor Ulitsky, Ron Shamir, and Martin Kupiec. Different sets of qtls influence fitness variation in yeast. Molecular systems biology, 6(1):346, 2010.

44. Brant Weinstein and Frank Solomon. Phenotypic consequences of tubulin overproduction in saccharomyces cerevisiae: differences between alpha-tubulin and beta-tubulin. Molecular and cellular biology, 10(10):5295–5304, 1990.

45. Haoping Liu, Janet Krizek, and Anthony Bretscher. Construction of a gal1-regulated yeast cdna expression library and its application to the identification of genes whose overexpression causes lethality in yeast. Genetics, 132(3):665–673, 1992.

46. Koji Makanae, Reiko Kintaka, Takashi Makino, Hiroaki Kitano, and Hisao Moriya. Identification of dosage-sensitive genes in saccharomyces cerevisiae using the genetic tug-of-war method. Genome research, 23(2):300–311, 2013.

47. DeElegant Robinson, Michael Place, James Hose, Adam Jochem, and Audrey P Gasch. Natural variation in the consequences of gene overexpression and its implications for evolutionary trajectories. Elife, 10:e70564, 2021.

48. Nathaniel P Sharp, Linnea Sandell, Christopher G James, and Sarah P Otto. The genomewide rate and spectrum of spontaneous mutations differ between haploid and diploid yeast. Proceedings of the National Academy of Sciences, 115(22):E5046–E5055, 2018.

49. Kaitlin J Fisher, Sean W Buskirk, Ryan C Vignogna, Daniel A Marad, and Gregory I Lang. Adaptive genome duplication affects patterns of molecular evolution in saccharomyces cerevisiae. PLoS genetics, 14(5):e1007396, 2018.

50. Enikö Zörgö, Karolina Chwialkowska, Arne B Gjuvsland, Elena Garré, Per Sunnerhagen, Gianni Liti, Anders Blomberg, Stig W Omholt, and Jonas Warringer. Ancient evolutionary trade-offs between yeast ploidy states. PLoS genetics, 9(3):e1003388, 2013.

51. John H Gillespie. Molecular evolution over the mutational landscape. Evolution, pages 1116– 1129, 1984.

52. H Allen Orr. The distribution of fitness effects among beneficial mutations. Genetics, 163(4):1519–1526, 2003.

53. Julia Muenzner, Pauline Trebulle, Federica Agostini, Christoph B Messner, Martin Steger, Andrea Lehmann, Elodie Caudal, Anna-Sophia Egger, Fatma Amari, Natalie Barthel, et al. The natural diversity of the yeast proteome reveals chromosome-wide dosage compensation in aneuploids. bioRxiv, 2022.

54. James Hose, Leah E Escalante, Katie J Clowers, H Auguste Dutcher, DeElegant Robinson, Venera Bouriakov, Joshua J Coon, Evgenia Shishkova, and Audrey P Gasch. The genetic basis of aneuploidy tolerance in wild yeast. Elife, 9:e52063, 2020.

55. Jackson Peter, Matteo De Chiara, Anne Friedrich, Jia-Xing Yue, David Pflieger, Anders Bergström, Anastasie Sigwalt, Benjamin Barre, Kelle Freel, Agnès Llored, et al. Genome evolution across 1,011 saccharomyces cerevisiae isolates. Nature, 556(7701):339–344, 2018.

56. Romain Koszul, Bernard Dujon, and Gilles Fischer. Stability of large segmental duplications in the yeast genome. Genetics, 172(4):2211–2222, 2006.

57. Ana B Oromendia, Stacie E Dodgson, and Angelika Amon. Aneuploidy causes proteotoxic stress in yeast. Genes & development, 26(24):2696–2708, 2012.

58. Luke Hakes, John W Pinney, Simon C Lovell, Stephen G Oliver, and David L Robertson. All duplicates are not equal: the difference between small-scale and genome duplication. Genome biology, 8(10):1–13, 2007.

59. Andreas Wagner. Energy constraints on the evolution of gene expression. Molecular biology and evolution, 22(6):1365–1374, 2005.

60. Reiner A Veitia, Samuel Bottani, and James A Birchler. Gene dosage effects: nonlinearities, genetic interactions, and dosage compensation. Trends in Genetics, 29(7):385–393, 2013.

61. Matthew Scott, Carl W Gunderson, Eduard M Mateescu, Zhongge Zhang, and Terence Hwa. Interdependence of cell growth and gene expression: origins and consequences. Science, 330(6007):1099–1102, 2010.

62. Eyal Metzl-Raz, Moshe Kafri, Gilad Yaakov, Ilya Soifer, Yonat Gurvich, and Naama Barkai. Principles of cellular resource allocation revealed by condition-dependent proteome profiling. eLife, 6, 2017.

63. Timothy R Hughes, Matthew J Marton, Allan R Jones, Christopher J Roberts, Roland Stoughton, Christopher D Armour, Holly A Bennett, Ernest Coffey, Hongyue Dai, Yudong D He, et al. Functional discovery via a compendium of expression profiles. Cell, 102(1):109–126, 2000.

64. Anjali Mahilkar, Namratha Raj, Sharvari Kemkar, and Supreet Saini. Selection in a growing colony biases results of mutation accumulation experiments. Scientific Reports, 12(1):1–12, 2022.

65. Lindi M Wahl and Deepa Agashe. Selection bias in mutation accumulation. Evolution, 76(3):528–540, 2022.

66. Neil Savage. Bioinformatics: big data versus the big c. Nature, 509(7502):S66–S67, 2014.

67. Jonathan D Groves, Pierre Falson, Marc Le Maire, and MJ Tanner. Functional cell surface expression of the anion transport domain of human red cell band 3 (ae1) in the yeast saccharomyces cerevisiae. Proceedings of the National Academy of Sciences, 93(22):12245–12250, 1996.

68. Martin A Nowak, Natalia L Komarova, Anirvan Sengupta, Prasad V Jallepalli, Ie-Ming Shih, Bert Vogelstein, and Christoph Lengauer. The role of chromosomal instability in tumor initiation. Proceedings of the National Academy of Sciences, 99(25):16226–16231, 2002.

69. Yoh Iwasa, Franziska Michor, and Martin A Nowak. Stochastic tunnels in evolutionary dynamics. Genetics, 166(3):1571–1579, 2004.

70. Michael Lynch and Adam Abegg. The rate of establishment of complex adaptations. Molecular biology and evolution, 27(6):1404–1414, 2010.

71. Michael Lynch. Scaling expectations for the time to establishment of complex adaptations. Proceedings of the National Academy of Sciences, 107(38):16577–16582, 2010.

72. Brigida Gallone, Jan Steensels, Troels Prahl, Leah Soriaga, Veerle Saels, Beatriz Herrera-Malaver, Adriaan Merlevede, Miguel Roncoroni, Karin Voordeckers, Loren Miraglia, et al. Domestication and divergence of saccharomyces cerevisiae beer yeasts. Cell, 166(6):1397–1410, 2016.

73. Shou-Fu Duan, Pei-Jie Han, Qi-Ming Wang, Wan-Qiu Liu, Jun-Yan Shi, Kuan Li, Xiao-Ling Zhang, and Feng-Yan Bai. The origin and adaptive evolution of domesticated populations of yeast from far east asia. Nature communications, 9(1):1–13, 2018.

74. Yuan O Zhu, Gavin Sherlock, and Dmitri A Petrov. Whole genome analysis of 132 clinical saccharomyces cerevisiae strains reveals extensive ploidy variation. G3: Genes, Genomes, Genetics, 6(8):2421–2434, 2016.

75. Katy C Kao, Katja Schwartz, and Gavin Sherlock. A genome-wide analysis reveals no nuclear dobzhansky-muller pairs of determinants of speciation between s. cerevisiae and s. paradoxus, but suggests more complex incompatibilities. PLoS genetics, 6(7):e1001038, 2010.

76. Anna M Selmecki, Yosef E Maruvka, Phillip A Richmond, Marie Guillet, Noam Shoresh, Amber L Sorenson, Subhajyoti De, Roy Kishony, Franziska Michor, Robin Dowell, et al. Polyploidy can drive rapid adaptation in yeast. Nature, 519(7543):349–352, 2015.

77. Mia Jaffe, Gavin Sherlock, and Sasha F Levy. iseq: a new double-barcode method for detecting dynamic genetic interactions in yeast. G3: Genes, Genomes, Genetics, 7(1):143–153, 2017.

## References

[1] Kimura M. On the probability of fixation of mutant genes in a population. Genetics. 1962 Jun;47(6):713.

[2] Gerrish PJ, Lenski RE. The fate of competing beneficial mutations in an asexual population. Genetics. 1998 Mar;102:127-44.

[3] Schiffels S, SzöllHosi GJ, Mustonen V, Lässig M. Emergent neutrality in adaptive asexual evolution. Genetics. 2011 Dec 1;189(4):1361–75.

[4] Haldane JB. A mathematical theory of natural and artificial selection, part V: selection and mutation. In Mathematical Proceedings of the Cambridge Philosophical Society 1927;23(7):838–844). Cambridge University Press.

[5] Yona AH, Manor YS, Herbst RH, Romano GH, Mitchell A, Kupiec M, Pilpel Y, Dahan O. Chromosomal duplication is a transient evolutionary solution to stress. Proceedings of the National Academy of Sciences. 2012 Dec 18;109(51):21010–5.

[6] Romano GH, Gurvich Y, Lavi O, Ulitsky I, Shamir R, Kupiec M. Different sets of QTLs influence fitness variation in yeast. Molecular systems biology. 2010;6(1):346.

